# Mating imperatives drive plasticity of the daily temporal niche in fruit flies

**DOI:** 10.64898/2026.07.02.736183

**Authors:** Sagnik Ghosh, Ping Zhong, Caroline Suray, Junaid Mir, Tingting Chen, Antonio Palazzo, Vincent Rincheval, François Rouyer, Abhishek Chatterjee

## Abstract

Animals confine their daily activity to distinct diurnal, crepuscular, or nocturnal windows. It remains unclear how hundreds of sympatric species sharing the same temporal niche manage to coexist. To address this, we monitored the daily activity of the twilight-active, species-rich Drosophila genus under naturalistic conditions. Here, we show that intense sociosexual interactions rapidly drive a species-specific reformatting of their canonical crepuscular niche. The dominant sensory modality used for sexual communication predicts niche shift direction: reliance on chemosensation for courtship redirects behavioral activity into the night, while visual reliance shifts it into the day. This temporal plasticity bypasses the circadian clock, instead operating via a conserved dopaminergic pathway that simultaneously inhibits sleep and sustains sexual motivation. Our results reveal how mating imperatives enable conditional colonization of otherwise restricted temporal windows. Ultimately, by driving the divergence of previously overlapping niches, sociosexually induced temporal plasticity provides a powerful mechanism for sympatric coexistence in crowded environments.

## INTRODUCTION

The environment in which animals live undergoes dramatic daily cycles as a direct consequence of Earth’s rotational motion. Animals deploy a combination of anticipatory adaptations and reactive responses to cope with the periodic changes in their ambient environment. This entails restriction of behavioral activity to specific phases of the solar day^1–3^. The resulting species-specific timing of locomotor activity operationally defines an animal’s daily temporal niche^4,5^. Partitioning of the temporal niche among closely related sympatric species reduces competition and paves the way for rapid evolutionary diversification during adaptive radiations^5^.

Under naturalistic settings, the notion of a rigid, species-specific temporal niche is challenged. Nocturnal rodents facing hunger or cold, for example, gradually switch to diurnality to reduce thermo-energetic expenditure^4,6,7^. In endothermic mammals, metabolic demand can drive this switch by resetting the circadian clockwork through defined energy-sensing pathway^8^. In contrast, being physiologically tethered to ambient temperature and humidity conditions, ectothermic insects face a distinct set of constraints. Do these ectotherms lack such active flexibility, or do they switch temporal niche following fundamentally different eco-physiological imperatives?

*Drosophila* flies are thought to be crepuscular, restricting activity to dawn and dusk^9,10^. This bimodal activity pattern persists across seasons, except on excessively hot days, when thermal stress triggers escape behavior^11,12^. Alongside these abiotic factors, potent biotic variables like the social environment further fine-tune daily activity rhythms under constant laboratory conditions^13,14^. We here demonstrate that in drosophilids, sociosexual interactions drive rapid remodeling of the temporal niche under naturalistic outdoor settings. While the clock and abiotic pressures establish a static temporal scaffold, sociosexual drive overlays an additional degree of freedom, allowing conditional niche expansion. Our findings suggest that in the ephemeral environment where insects thrive, dopaminergic sexual incentive generates the temporal partitioning necessary for species coexistence.

## RESULTS

### *Drosophila* temporal niche is reshaped under sociosexual settings

At the end of summer, when overripe fruits begin to ferment, wild *D. melanogaster* gather in dense aggregates to feed, mate, and lay eggs^15,16^. Using high-resolution actigraphic monitors, we asked when, during the 24-h cycle, groups of flies become behaviorally active under semi-natural conditions (Extended Data Fig. 1a). Mixed-sex groups displayed striking nocturnality in European summer nights (Fig. 1a-b and Extended Data Fig. 1b). Warmer night temperatures rather than photoperiod facilitated the nocturnal switch (Extended Data Fig. 1c-f).

**Figure 1:**
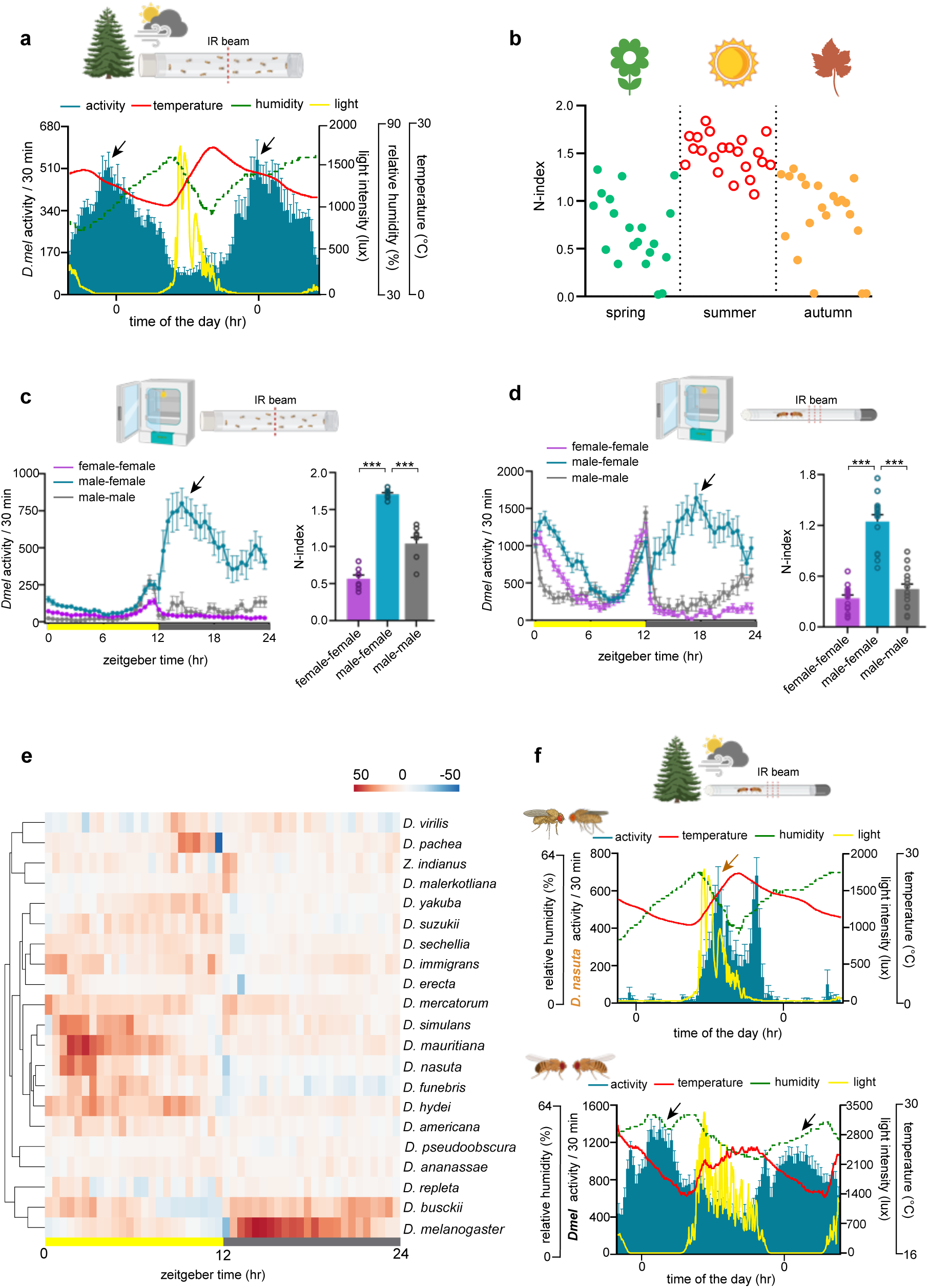
Flies under sociosexual settings switch their temporal niches. **(a)** Locomotor activity profile of *D. melanogaster* mixed-sex groups of flies reveals pronounced nocturnal activity (black arrow) under seminatural summer conditions in Versailles, France (n=8, 30 flies per group). On the x-axis, 0 corresponds to midnight. **(b)** Seasonal variation in the nocturnal activity of fly groups under seminatural conditions (n=8, 30 flies per group). Datapoints are presented in temporal order. **(c-d)** (left) Locomotor activity profiles of *Dmel* flies reveal prominent night peaks (black arrow) in mixed-sex groups (c), as well as pairs (d) under laboratory conditions (12h:12h light-dark cycle, constant 25°C; n=8 for groups of 30 flies; n=15 for pairs). (right) Quantification of the night peak (N-index). N-index reflects the relative concentration of activity at night (see Methods for details). **(e)** Heatmap along with cluster analysis illustrating the time-dependent relative difference in locomotor activity of mixed-sex pairs compared to individual males across 21 Drosophila species under laboratory conditions (n=14). Color scale reflects differences in normalized locomotor activity. **(f)** Under seminatural conditions, activity profiles of mixed-sex pairs of *D. nasuta* (top) and *D. melanogaster* (bottom) flies are marked by a day peak (copper arrow) and a night peak (black arrow), respectively (n=14 per species). Data are presented as mean ± SEM. Statistical comparisons were performed using a two-tailed unpaired t-test. Significance levels: *p<0.05; **p<0.01; *** p<0.001. See Supplementary Table 1 for details on statistics. Schematics were created using BioRender.

Mixed-sex groups remained nocturnal even when environmental fluctuations were reduced to square-wave light cycles in laboratory settings (Fig. 1c). Their nocturnal activity eclipsed the well-documented morning and evening peaks, which in contrast, continued to dominate the activity profile of the same-sex groups (Fig. 1c). The nocturnal activity peak emerged robustly across assay formats and detection methods at population densities typical of natural aggregates (Supplementary Video 1 and Extended Data Fig. 2a-b)^17^. Even a pair of male and female, though not same-sex pairs, produced a distinct night peak (Fig. 1d). Sociosexual opportunity rapidly and reversibly induced this temporal plasticity (Extended Data Fig. 2c). The night peak was built by a synchronous rise in male and female activity associated with intense sexual interaction (Supplementary Video 2), leading to an upswing of the copulation rhythm toward the end of the night (Extended Data Fig. 2d-f). Thus, a unique behavioral state, fundamentally distinct from that underlying the canonical morning and evening peaks, marked the night peak (Extended Data Fig. 2g, Supplementary Table 4). The night peak did not merely mirror the rhythm in close-proximity encounters, which, in fact, peaked after daybreak (Extended Data Fig. 2h-i)^18,19^. Moreover, of the so-called proximity encounters, barely half reflected courtship (Extended Data Fig. 2j)^20^.

We next asked whether sociosexual interactions may promote temporal niche partitioning across *Drosophila* species that are crepuscular as individuals (Extended Data Fig. 3a). In mixed-sex pairs, the activity phase of different species stably shifted in opposite directions (Fig. 1e and Extended Data Fig. 3a-b). The shift towards nocturnality was most prominent in distantly related *D. melanogaster* and *D. busckii*, suggestive of repeated, independent evolution of night-shift capacity (Fig. 1e and Extended Data Fig. 3b-c). In contrast, the diurnal shift was most noticeable in *D. nasuta*, *D. simulans,* and *D. mauritiana*. Several others, such as *D. ananassae* and *D. pseudoobscura*, remained crepuscular (Fig. 1e and Extended Data Fig. 3a-b). These species-specific activity-phase shifts were also evident under seminatural outdoor conditions (Fig. 1f). Among the four species of the melanogaster subgroup (Extended Data Fig. 3c), *D. melanogaster* alone exhibited a nocturnal shift, while the others showed varying degrees of diurnal shift, consistent with a *D. melanogaster*-specific character displacement. These cosmopolitan *D. melanogaster* flies occupy overlapping niches with many other drosophilids, including *D. nasuta*, *D. ananassae*, and *D. simulans*^21,22^. To test whether our findings of activity phase-shifts observed in intraspecific interactions translate to the community level, we assembled mixed-sex groups of these naturally sympatric pairs – *Dmel*-*Dnas*, *Dmel*-*Dana*, and *Dmel*-*Dsim*, some of which can even produce sterile hybrids^23^. Species-specific activity-phase shifts indicative of temporal niche partitioning were again evident in these model assemblages, where individuals could interact both within and across species (Extended Data Fig. 3d-f).

### Distinct sensory determinants of diurnal and nocturnal shifts

The contrasting directions of the niche-shifts suggest that species differ in the mechanisms that couple sociosexual activity to specific phases of the day-night cycle. Species-specific temporal niche shifts were mirrored by corresponding biases in courtship timing: night-shifting species courted more at night, whereas day-shifting species showed higher courtship during the day (Fig. 2a). With increasing daylight intensity, night-shifting species showed diminishing shift to nocturnality, while the day-shifting species had an enhanced diurnal shift (Fig. 2b and Extended Data Fig. 4a-b). Indeed, intense light enhances male courtship in otherwise night-shifting *D. melanogaster*, facilitating visually-guided tracking of the females during daytime^24,25^. Given the contrasting effects of light, we asked if the relative importance of sensory modalities that shape niche-shift direction differs between these two groups. The day-shifting species invest more in visual processing, with relatively enlarged optic lobes, while night-shifting species prioritize non-photic sensory processing (Fig. 2c).

**Figure 2:**
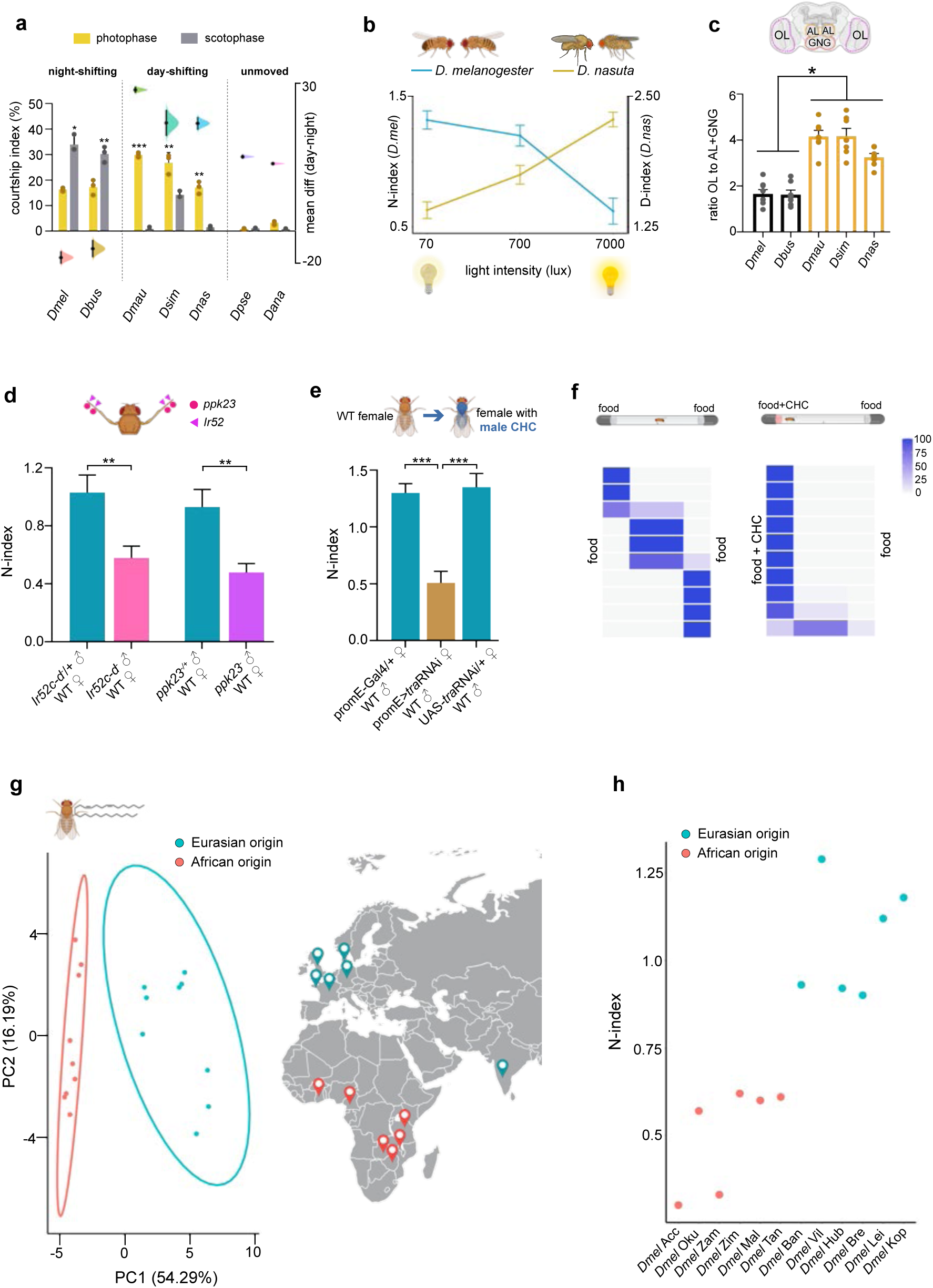
Divergence in sensory cues shapes temporal niche shifts. **(a)** Courtship indices in the photophase and scotophase (bars) and their mean difference (dots with gaussian ribbons) across different *Drosophila* species (n=10, 3 repeats). Night-shifting species court more at night, while day-shifting species exhibit higher courtship during the day. **(b)** The variation of N and D-indices of *D. melanogaster* and *D. nasuta* pairs under increasing daylight intensity (n=14 for each). The two species exhibit contrasting effects of daylight on their sociosexual peak. **(c)** Relative sensory allocation, shown as the ratio of optic lobe (OL) to antennal lobe (AL) and gnathal ganglion (GNG), in the males of day and night-shifting *Drosophila* species (n=7 brains per species). **(d)** Male-specific knockout of *Ir52c-d* and *ppk23* receptors suppresses the night peak (n=14 for both). WT refers to wild-type. **(e)** Modulation of night peak due to masculinization of female CHC profiles via *tra*-RNAi (n=14). **(f)** Positional heatmap showing individual males’ spatial preference when given a choice between food (left) and food spiked with female CHCs (right) (n=10). **(g)** Principal Component Analysis (left) on the female CHC profiles of wild-caught *Dmel* flies reveals two distinct groups: Eurasian and African (right). **(h)** K-means clustering (explained variance: 79%) separates the N-indices of the wild-caught *Dmel* pairs based on their geographical origins. Data are presented as mean ± SEM. Statistical comparisons were performed using a two-tailed unpaired t-test and one-way ANOVA. Significance levels: *p<0.05; **p<0.01; *** p<0.001. See Supplementary Table 1 and Supplementary Table 3 for details on statistics and genetic manipulations, respectively. Schematics were created using BioRender.

Which non-visual modality guides niche-shift toward nocturnality? Screening of *D. melanogaster* sensory mutants revealed that disruption of *piezo*-mediated mechanosensation led to a modest reduction of nocturnality, whereas the loss of taste (*Poxn* mutant) entirely abolished the night peak (Extended Data Fig. 4c). This reliance on taste mapped to the cuticular pheromone receptors *Ir52* and *ppk23*^26,27^ in the foreleg tarsi of males but not females (Fig. 2d and Extended Data Fig. 4d-f). Removal of male foreleg tarsi in the night-shifting *D. busckii* also prevented niche-shift (Extended Data Fig. 4g). Consistent with the divergence in sensory bias, the temporal plasticity of the day-shifting species was recalcitrant to tarsi removal (Extended Data Fig. 4g).

Tarsal chemoreceptors process cuticular hydrocarbons (CHCs) that signal sex, species identity, and individual quality, providing a balance of stimulatory and inhibitory cues during courtship^26–30^. Consistent with this, *Dmel* males did not produce the night peak with heterospecific females, unless their cuticular hydrocarbon (CHC) profile contained *Dmel* female-specific dienes (Extended Data Fig. 5a)^31^. On the other hand, masculinizing the sexually dimorphic CHCs of conspecific females diminished the night peak. Thus, sociosexual signaling via female CHCs supports the nocturnal niche-shift in *Dmel* flies (Fig. 2e). We then asked if the loss of night peak upon disruption of CHC reception is a mere consequence of courtship suppression. Reception of the sexually dimorphic CHCs was largely dispensable for courtship and mating (Extended Data Fig. 5b)^28,32^. Instead, CHCs served as a general elicitor of male attraction (Fig. 2f and Extended Data Fig. 5c), consistent with their role as aphrodisiac^33^.

In light of our finding that male detection of female CHCs drives nocturnal shifts, we asked whether naturally varying female CHC profile drives intraspecies variations in the night peak. GC-MS revealed across-the-board divergence between the CHC profiles of wild-caught sub-Saharan and Eurasian *Dmel* females (Fig. 2g and Extended Data Fig. 5d, Extended Data Table 1)^34^. Their night peaks also segregated by geography, suggestive of its association with the female CHC profile (Fig. 2h and Extended Data Fig. 5e-f). The Eurasian strains showed higher night peaks in general (Fig. 2h and Extended Data Fig. 5e). Oenocyte-depleted CHC-less females treated with Eurasian CHC extract elicited a higher nocturnal peak than those carrying African CHCs, establishing causality (Extended Data Fig. 5g). Collectively, these results reveal that the sensory bias embedded in the nervous system defines the direction of niche change, while the potency of sociosexual signals scales the extent of that temporal displacement.

### A shared dopaminergic mechanism instructs temporal niche plasticity

Circadian clock-generated morning and evening peaks of locomotor activity persist under constant external environments. However, the night peak was not distinguishable in prolonged constant darkness (DD), under which the MF pairs gradually became arrhythmic (Extended Data Fig. 6a-c). This prompted us to test whether the night peak is modified in mixed-sex pairs of clock mutants under light– dark conditions. Clock-less *per*⁰ and *tim*⁰ flies retained the night peak (Fig. 3a-b and Extended Data Fig. 6d-e), while their morning and evening anticipatory peaks were absent. Dispensability of the circadian system for night peak indicates that sociosexually evoked niche-shift can materialize without an endogenous clockwork.

**Figure 3:**
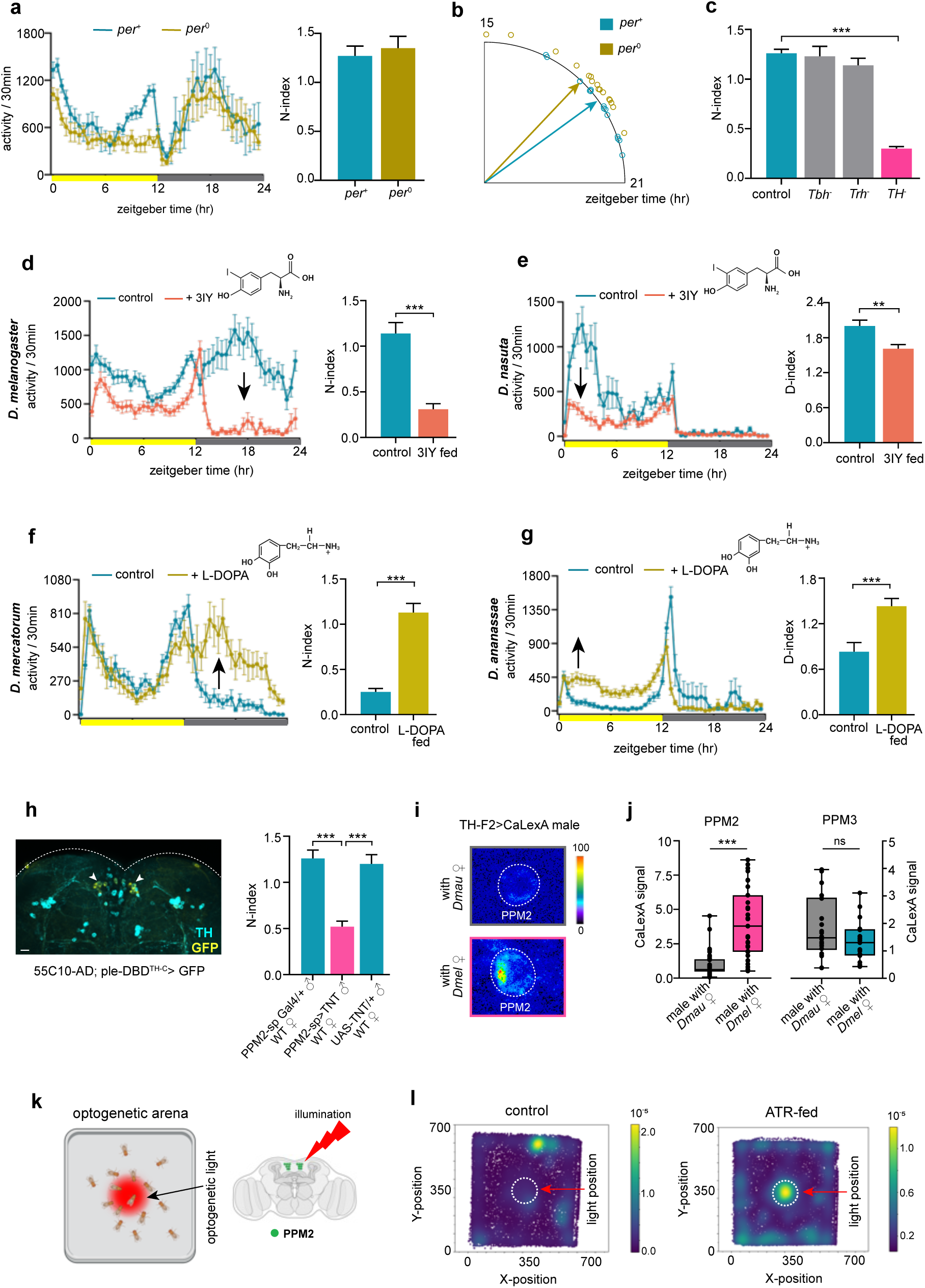
Dopamine mediates divergent temporal niche shifts. **(a)** 24-hour locomotor activity profile (left) and N-index (right) of the mixed-sex pairs of circadian clock mutant (*per*^0^) and control flies (n=14). **(b)** The phase of the night peak does not differ between *per*^0^ and control pairs (n=14). **(c)** Night peak of flies deficient in octopamine (*Tbh^-^*), serotonin (*Trh^-^*), and dopamine (*TH^-^*) (n=14 for each). **(d-g)** Effect of pharmacological manipulation of dopamine signalling on the activity profile and N-or D-index. *D. melanogaster* (d) and *D. nasuta* (e) pairs fed with the antagonist 3IY decreased their sociosexual night and day activity, respectively. *D. mercatorum* (f) and *D. ananassae* pairs (g) fed with the agonist L-DOPA increased their night and day activity, respectively (n=16 per species). **(h)** (left) PPM2 dopaminergic neurons in the *Dmel* brain addressed by a split-Gal4 (55C10∩ple^TH-C^). (right) Suppression of the night peak by tetanus toxin (TNT)-mediated silencing of the male PPM2 neurons (55C10∩ple^TH-C^>TNT, tsh-Gal80) (n=14) **(i)** Calcium levels in PPM2 neurons of *Dmel* males after interaction with either *Dmau* (top) or *Dmel* (bottom) females. **(j)** Quantification of the calcium (CaLexA) signal in PPM2 and PPM3 neurons of *Dmel* males under the same conditions (n≥7 brains). **(k)** Schematics of the optogenetic experimental setup for PPM2 activation. **(l)** During optogenetic stimulation, all-trans-retinal (ATR) fed 55C10∩ple^TH-C^>CsChrimson males proactively spent time in the illumination zone (right), unlike control males (left) (n=12). Color scales in the spatial heatmap indicate positional occupancy frequencies. White circles indicate the optogenetic illumination zone. Scale bar is 10 µm. Bar graphs show mean ± SEM. In box plots, horizontal lines represent the median, box bounds indicate the 25th and 75th percentiles, and whiskers extend from minimum to maximum values. Statistical comparisons were performed using a two-tailed unpaired t-test. Significance levels: *p<0.05; **p<0.01; *** p<0.001. See Supplementary Table 1 and Supplementary Table 3 for details on statistics and genetic manipulations, respectively. Schematics were created using BioRender.

As biogenic amines are critical regulators of behavioral plasticity across phylogeny^35^, we reasoned, they could be key to the niche-shift capacity. Mutant pairs deficient in the biosynthesis of neuronal dopamine, but not the other monoamines, lacked the night peak (Fig. 3c). Consistent with this, the night peak was also absent in transgenic flies whose dopamine neurons in the brain were silenced with tetanus toxin (DAT>TNT, tsh-Gal80) as well as in flies subjected to adult-specific silencing of these neurons via conditional expression of the hyperpolarizing potassium channel Kir2.1 (DAT>Kir, tub-Gal80^TS^) (Extended Data Fig. 7a). Of the four known dopamine receptors in *Drosophila*, the D1-like Dop1R2 was necessary (Extended Data Fig. 7b). Consistent with this, the night peak could be rescued in the dopamine-deficient mutant fed with the precursor of dopamine, *i.e.*, L-DOPA/carbidopa^36^, but not in the mutant lacking Dop1R2 (Extended Data Fig. 7c). We then administered the dopamine synthesis inhibitor 3-Iodo-L-tyrosine (3IY)^37^ to different *Drosophila* species exhibiting either day or night-specific sociosexual activity peaks. 3IY treatment effectively suppressed the niche-shift capacity of both groups while leaving circadian clock-driven activity peaks at morning and evening intact (Fig. 3d-e and Extended Data Fig. 7d-f). On the other hand, L-DOPA/carbidopa supplementation enhanced the niche-shift in *D. mercatorum*, which otherwise showed limited temporal plasticity (Fig. 3f). Similarly, *D. ananassae* and *D. pseudoobscura*, species that typically did not alter their daily activity patterns in a sociosexual context, could also exhibit a time-specific activity increase when fed dopamine agonists (Fig. 3g and Extended Data Fig. 7g). These findings suggest that dopamine signaling is a common mediator of temporal niche plasticity across several Drosophila species.

To pinpoint the specific group of dopamine neurons (DANs) driving niche shift in *Dmel* brain, we used progressively restrictive Gal4 lines targeting subsets of ∼250 DANs, organized into ten clusters^38^. Constitutive synaptic inhibition (via TNT), as well as adult-specific electrical silencing (via Kir2.1) of the PPM2 cluster, which contains 7-8 DANs, abolished the night peak (Fig. 3h and Extended Data Fig. 7h-i). In contrast, transmission from PPM3 DANs, which upholds circadian (DD) locomotor rhythms in isolated individuals^39^, was dispensable (Extended Data Fig. 7j).

Prolonged conspecific sociosexual interaction drove sustained activation of male PPM2 neurons, revealed by increased CaLexA signal, a marker of cumulative neural activity^40^ (Fig. 3i-j). Neighbouring PPM3 DANs remained unaltered, confirming that sustained activation is selective to PPM2 (Fig. 3j). Thus, persistent sociosexual activation of male PPM2 DANs triggers nocturnality.

Although inhibition of PPM2 neurons suppressed the night peak, it did not alter acute courtship toward virgin females on short timescales (Extended Data Fig. 7k). Which behavioral process, then, is regulated by the PPM2 DANs? To answer this, we optogenetically activated them in single males (55C10∩ple^TH-C^>CsChrimson), which resulted in a marked change in their motivational state. Single males actively sought and remained within the illuminated region of the arena where PPM2 neurons could be proactively stimulated (Fig. 3k-l and Extended Data Fig. 7l). This self-directed occupancy of the stimulation zone indicates that PPM2 activity conveys an intrinsically reinforcing signal. Other DANs involved in appetitive reinforcement were not required for night peak generation, underscoring the unique role of PPM2 neurons in converting sociosexual experience into enduring changes in daily activity patterns (Extended Data Fig. 7j).

### The neural circuit for night peak in *D. melanogaster*

Mated *Dmel* mixed-sex pairs produced a robust night peak night after night (Fig. 4a), even though most courtship attempts were unfruitful. Positively valenced memory of prior mating^41^ did not drive this persistence, as learning mutants showed normal night peaks (Extended Data Fig. 8a). Nor could it be explained by steadily mounting sexual motivation in unsatiated males, as ∼95% pairs remated at least once during two days of cohabitation in our assay system. However, forced ejaculation (Crz>NaChBac), which oversatiates males^41,42^, extinguished the peak, as did hyper-receptive *sex-peptide receptor* (SPR) mutant females (Fig. 4b and Extended Data Fig. 8b). Conversely, non-receptive *fru*^M^ females and nearly asexual *fru*^F^ males suppressed the night peak (Fig. 4c and Extended Data Fig. 8c). Thus, the night peak is underpinned by a calibrated equilibrium of sexual motivation between the two sexes.

**Figure 4:**
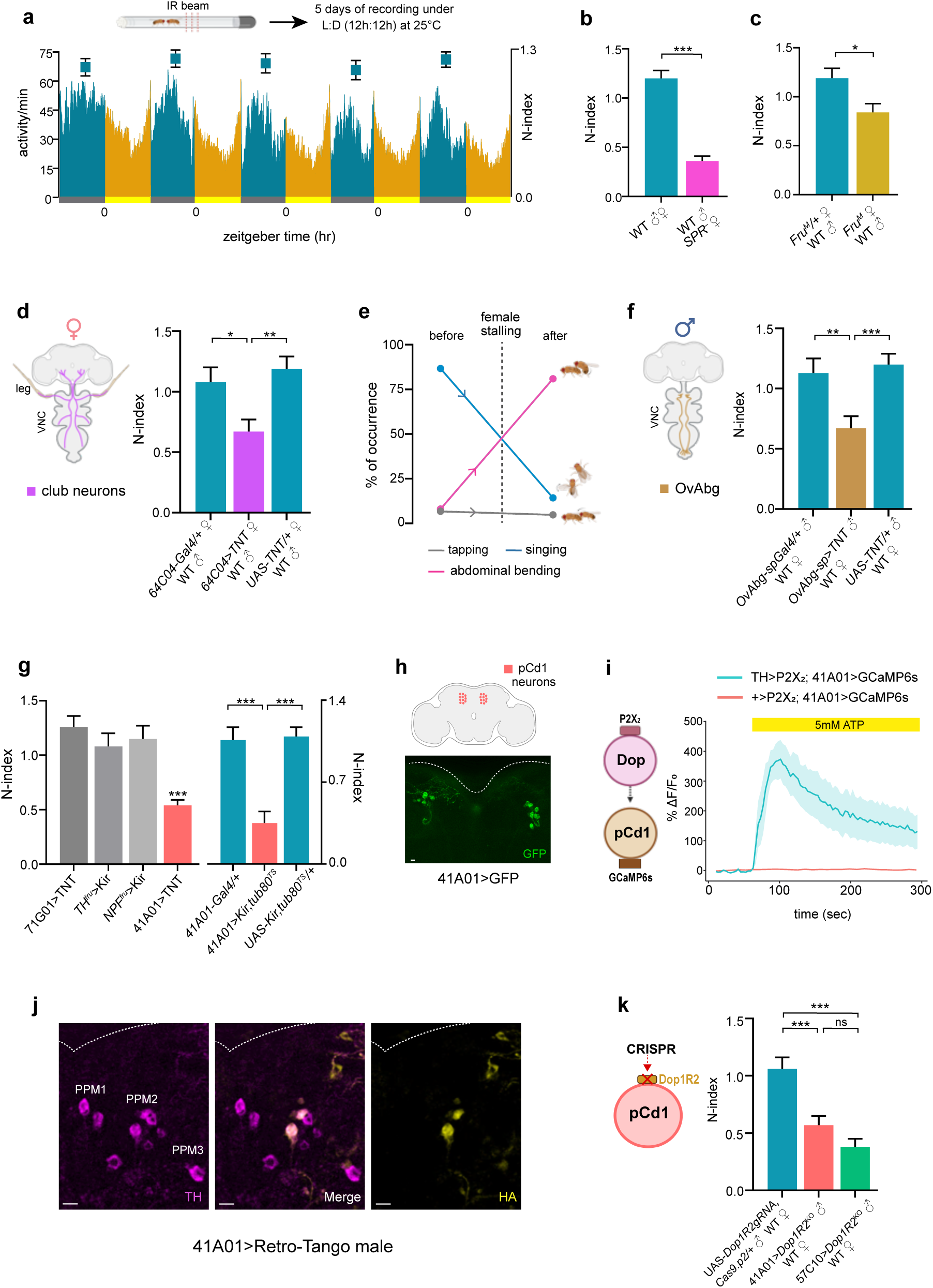
PPM2–pCd1 connection regenerates sex drive in *D. melanogaster*. **(a)** Locomotor activity rhythms and N-indices (squares above) of mixed-sex *Dmel* pairs displaying the night peak over five consecutive days (n=16). **(b-c)** Effects of change in female receptivity on the night peak: (b) knockout of the sex-peptide receptor (SPR), and (c) misexpression of the male-specific *fruitless* isoform (*fru*^4–40^*/fru*^M^) in females, both reduce the night peak (N=14 for both). **(d)** (left) Club neurons that regulate female-stalling. (right) Female-specific silencing of the club neurons (64C04>TNT) reduces the night peak (n=14). **(e)** Changes in the occurrence of different courtship subroutines displayed by the males before and after female-stalling. **(f)** (left) OvAbg neurons that regulate male abdominal bending. (right) Reduction of the night peak due to TNT-mediated silencing of OvAbg neurons (VT038154∩dsx>TNT) in males (n=14). **(g)** (left) Influence of silencing different nodes of the courtship circuit (P1/pC1, TH∩fru, NPF∩fru, and pCd1) on the night peak (n=14 for each). (right) Adult-specific conditional silencing of pCd1 neurons, induced by temperature-dependent Gal80 inactivation and subsequent Gal4-driven Kir2.1 expression, suppresses the night peak (n=14). **(h)** pCd1 neurons in the male brain. Identical labeling patterns were observed across 3 independent brains. **(i)** Calcium (GCaMP6s) transients in the pCd1 neurons (41A01-Gal4) of a representative male brain in response to P2X_2_-mediated activation of the dopamine neurons (*ple*^TH^-LexA). **(j)** Retro-synaptic tracing from the male pCd1 neurons identifies PPM2 (co-stained with TH antibody) as their upstream partner. Results were consistently reproduced across 5 independent brain preparations. **(k)** Night peak is suppressed by CRISPR knockout of Dop1R2 receptor in all neurons (57C10), or just pCd1 neurons (41A01) of males (n=14 for both). Scale bar is 10 µm. Data are presented as mean ± SEM. Significance levels: *p<0.05; **p<0.01; *** p<0.001. Statistical comparisons were performed using a two-tailed unpaired t-test and one-way ANOVA. See Supplementary Table 1 and Supplementary Table 3 for details on statistics and genetic manipulations, respectively. Schematics were created using BioRender.

Because copulation, the terminal indicator of female receptivity, is relatively infrequent, we reasoned that a more frequently expressed subtle female signal sustains the night peak. Female cue that does not inevitably lead to copulation is her transient stalling during mobile courtship^43^. Silencing the leg club neurons (64C04>TNT) that mediate this stalling^44^ diminished the night peak without altering her overall locomotor activity (Fig. 4d and Extended Data Fig. 8d). We noticed, female-stalling often promoted a transition in male behaviour from female-directed displays to abdominal bending, a self-directed consummatory action (Fig. 4e). Suppression of abdominal bending (OvAbg>TNT) abrogated the night peak (Fig. 4f). Repeated abdominal bending has been shown to gradually activate the PPM2 neurons^45^, linking it with the dopaminergic foundation of the night peak.

We next asked which node in the male courtship circuit is modulated by PPM2 (Extended Data Fig. 8e). Silencing the P1 command neurons (71G01>TNT, tsh-Gal80) or the *fru*⁺ dopaminergic and NPF neurons that feed into the P1 hub (NPF∩fru>Kir, TH∩fru>Kir), did not disrupt the night peak (Fig. 4g and Extended Data Fig. 8f). Indeed, P1-inactivated males can still display some sociosexual activity^46,47^. On the other hand, constitutively inhibiting neurotransmission (41A01>TNT, tsh-Gal80) and adult-specific electrical silencing (41A01>Kir, tub-Gal80^TS^) of a subset of *dsx*⁺/*fru*⁺ male pCd1 neurons markedly reduced the night peak (Fig. 4g-h and Extended Data Fig. 8f-g). pCd1 neurons sustain internal sociosexual states^48,49^, consistent with their requirement for maintaining the night peak. Physiological, anatomical, and behavioral analyses identified these neurons as the principal target of PPM2: dopaminergic activation elicited robust calcium rise in pCd1 neurons (Fig. 4i and Extended Data Fig. 8h); retro-synaptic tracing, corroborated by connectome mining, identified PPM2 as a direct upstream partner of pCd1 (Fig. 4j and Extended Data Fig. 8i); and CRISPR deletion of Dop1R2 from pCd1 disrupted the night peak (Fig. 4k and Extended Data Fig. 7b). These results place pCd1 as the dopamine-responsive persistence integrator that sustains the nocturnal uptick in sociosexual activity.

To sculpt the night peak, the PPM2-headed circuit must not only sustain sex-drive but also suppress night-time sleep. Prolonged sociosexual interaction markedly elevated calcium levels (69F08>CaLexA) in the sleep-debt-sensing R5 ring neurons of the ellipsoid body (Extended Data Fig. 9a)^50^. Yet its downstream dFB neurons, the effector arm of the sleep-homeostat (23E10>CaLexA), remained unchanged despite registration of the rising sleep pressure upstream in the circuit (Extended Data Fig. 9b). As it inhibits the dFB neurons^51,52^, PPM2 is poised to decouple the sensing of sleep debt from sleep execution during sociosexual interactions (Extended Data Fig. 9c). PPM2 silencing in male brains (55C10∩ple^TH-C^>TNT) elevated nighttime sleep in mixed-sex pairs, while leaving that of solitary males unaffected (Extended Data Fig. 9d-e). Thus, the sleep-regulatory role of PPM2 emerges specifically under a sociosexual context. Consistent with this PPM2-centric model, overactivation of these neurons in single males (55C10∩ple^TH-C^>NaChBac) generated a nocturnal shift (Extended Data Fig. 9f), whereas forced activation of the downstream dFB neurons in males attenuated this shift in mixed-sex pairs (Extended Data Fig. 9g). Therefore, to sculpt a nocturnal state of arousal, PPM2 engages a dual-channel circuit, *i.e.*, potentiation of sociosexual activity via pCd1 and suppression of sleep via dFB, bypassing the canonical homeostat flow (Fig. 5b).

**Figure 5:**
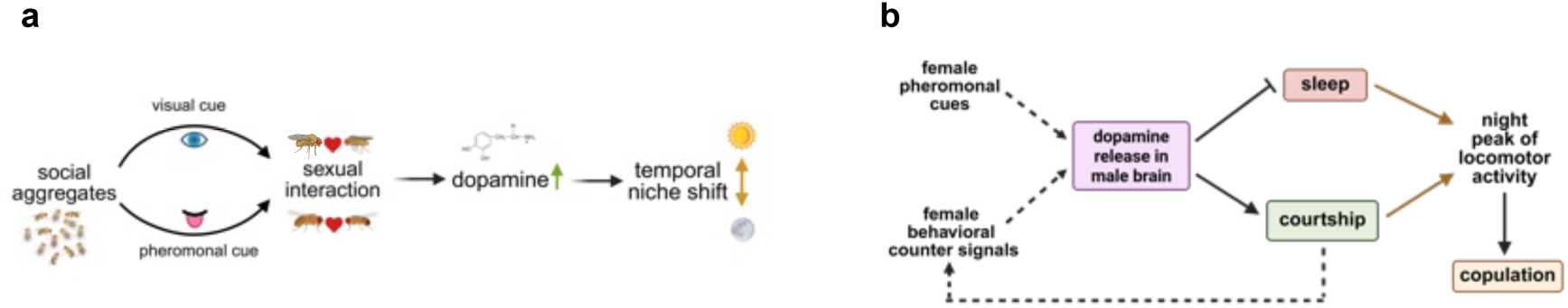
Mechanism underlying temporal niche-shifts. **(a)** Dopamine signaling drives temporal niche shifts across *Drosophila* species. Shifts towards daytime and nighttime are guided by visual and pheromonal cues, respectively. **(b)** In the male *Dmel* brain, dopamine neurons integrate pheromonal and behavioral inputs from female. Dopamine builds the night peak by suppressing sleep and promoting courtship. Schematics were created using BioRender.

## DISCUSSION

The intrinsic temporal niche of an organism is its daily window of activity evolved to cope with predictable abiotic oscillations. For the vinegar fly *Drosophila*, this results in a conserved crepuscular organization where plasticity is limited to minor phase adjustments—tracking of the seasonal shifts in dawn and dusk without altering the underlying activity structure^53–55^. Broader comparative studies have also confirmed that environmental fluctuations primarily scale the overall amount of locomotor activity of drosophilids, rather than driving a fundamental, sustained redistribution of the temporal niche^56^. However, studies carried out on isolated individuals under sterile laboratory conditions typically do not take into account the social dynamics of insect communities and the sensory complexities of their environment. Here, we show that density-dependent sociosexual interactions in *Drosophila* drive a rapid, reversible, and species-specific reformatting of the daily temporal niche under semi-natural outdoor conditions. Persistence of these niche shifts in community prototypes of sympatric drosophilids supports the temporal segregation of behavioral engagement across competing species. In evolutionary ecology, the traditional model for resource partitioning is morphological character displacement—famously exemplified by the beak size divergence of Darwin’s finches to partition trophic niche. Similarly, recent studies on African cichlid fishes have demonstrated that transitioning into a new temporal niche entails fixed evolutionary changes^5^. In contrast to such modification of the genomic hardware of organisms, sociosexually driven niche plasticity in *Drosophila* leverages neuromodulation to offer a rapidly reversible shortcut for species coexistence, suitable for insects thriving in highly dynamic microenvironments. Meanwhile, the underlying evolutionary divergence dictating the species-specific direction and extent of these shifts may represent a classic example of character displacement.

The circadian clock sculpts the temporal niche of individuals. The abiotic factors frequently leverage the circadian system to actuate temporal plasticity. For instance, nocturnal hyperactivity in *D. melanogaster* flies on moonlit nights or the emergence of an afternoon activity peak during extreme heat are both routed by the circadian system^10,11,55^. On the other hand, the biotic environment exerts limited influence on the core pacemaker of the vinegar flies^13,57,58^. Instead, it is the rhythmic output in overt behaviors that display prominent sensitivity to biotic cues like sociosexual interactions. For example, cohabiting mixed-sex pairs or groups of *D. melanogaster* flies display low-amplitude rest-activity cycles marked by pronounced sleep loss across both day and night^50,59–61^. While social cues reshape clock outputs, the pacemaker concurrently times sociosexual behaviors like close-proximity encounters that peak during the early morning hours under the dual guidance of light and clock^18,19^. Layered upon this clock-centric control of temporality, we demonstrate sociosexual interactions in *Drosophila* driving a powerful, clock-independent override that forces a dramatic shift in the temporal niche. A critical future direction will be to determine how this clock-independent mechanism interacts with clock-dictated daily proximity and copulation rhythms^18,19,62^. Even when a behavioral manifestation bypasses the core pacemaker, individual clock genes can still modulate rhythmic profiles through pleiotropic, non-circadian pathways. An elegant precedent is the *trans*-acting effect of *period* (*per*) 3′UTR splicing on the reverse-oriented *daywake* (*dyw*) gene, which recalibrates daytime sleep in response to changes in the abiotic environment, *i.e.*, ambient temperature, independent of functional PER protein^63^. A parallel clock-gene-linked, yet circadian-independent process could similarly gate the sociosexual induction of the night peak of locomotor activity in *D. melanogaster* flies. Furthermore, this capacity for an immediate, clock-independent behavioral override highlights a fundamental divergence in evolutionary strategy between insects and mammals. In mammals, within-lifetime temporal niche shifts are typically gradual, driven by persistent metabolic demands that slowly reset the circadian pacemaker^6^. This divergence likely reflects distinct functional requirements: while the circadian clock has evolved to buffer transient metabolic perturbations by integrating internal signals over longer timescales, such inertia is ill-suited for the reactive, volatile nature of sexual opportunity.

In nature, ecological succession, linked to the changing chemistry of fruit fermentation, operates along an extended timescale to shape the community composition of cohabiting *Drosophila* species^16,64^. Acting in parallel, species-specific shifts of the daily temporal niche, operating along a 24h timescale, should further constrain niche overlap within dense assemblages, mitigating interspecific competition and sterile hybridization. Even subtle differences in the timing of daily activity have been proposed to reinforce mating isolation in drosophilids^65^. While temporal niche plasticity is poised to minimize inter-species reproductive interference and optimize intra-species reproductive output, the physiological tax incurred by night-time sleep deprivation^66^ and male harassment likely opens up a hidden battleground for sexual conflict. This trade-off should effectively convert temporal niche plasticity into a double-edged sword. Physiological cost asymmetries between the sexes may arise from a male-specific neural hack, such as a dimorphic PPM2–dFB connection. Consequently, sleep loss during nocturnal courtship would leave a smaller footprint on the male nervous system. In contrast, females might perennially suffer from sleep deprivation and social jetlag.

In the *Drosophila* species (*e.g.*, *D. ananassae* and *D. pseudoobscura*) that did not modify their temporal niche under sociosexual context, females tend to remate only after several weeks^67^, while those modifying their crepuscular activity pattern were frequent rematers. Species like *D. nasuta, D. simulans,* and *D. mauritiana*, with sexually monomorphic cuticular hydrocarbons and light-dependent courtship involving postural displays^31,68^, turn more diurnal in mixed-sex settings. In contrast, those (*e.g.*, *D. melanogaster* and *D. busckii*) turning to nocturnality are able to court in the absence of light^31,69^, and carry sexually dimorphic cuticular pheromones that support conspecific mate recognition. Focusing on *D. melanogaster*, our results indicate that the shift to nocturnality is vetted by a bilayered sensory control that ensures the night peak is selectively deployed under conditions conducive to nocturnal mating. First, seasonally changing night temperatures open a permissive environmental gate. Subsequently, contact cues provide the acute trigger to drive a surge in sociosexual activity around midnight. Multimodal integration of contact-based chemical input from the broadly-tuned pheromone receptors *Ir52* and *ppk23* with Piezo-mediated mechanosensation enables the fly to continuously monitor the composition and density of its surrounding population. In cosmopolitan species, genetic variation in such sexual signaling pathways may support a cline in niche-change capacity, as higher latitudes restrict the reproductive window, increasing pressure for opportunistic niche expansion.

Our results suggest that the plasticity of the temporal niche is driven by a state-dependent dopamine-based mechanism across multiple *Drosophila* species. While divergence in sensory bias steers the direction and magnitude of these temporal shifts, the core engine remains a dopaminergic drive. Our working hypothesis (Fig. 5b) posits that PPM2 dopamine neurons in the *D. melanogaster* brain operate a homeostatic relaxation oscillator that tracks sex drive and sexual satiety. Around daybreak, successful copulation triggers a rapid discharge of the dopaminergic tone that encodes sex drive^70,71^. Consequently, during the forenoon, males exhibit low sexual motivation, which is further compounded by courtship conditioning^72^. At the same time, their innate attraction to the recently mated females remains inhibited by the anti-aphrodisiacs transferred to females during mating^73,74^. At nightfall, females gradually recovering from post-mating sex suppression^75^ and mate-guarding male manipulations^74^ begin to display cryptic receptive signals, despite remaining unreceptive to immediate copulation. As males detect female pheromonal cues and behavioral feedback, their courtship attempts intensify, replenishing the PPM2 dopaminergic tone^45^. As a result, the nocturnal activity peak is rebuilt through PPM2-mediated simultaneous inhibition of sleep and reinforcement of courtship. Taken together, the PPM2 neurons manage a behavior-guided daily cycle of reset and recovery of sex drive. This PPM2-headed circuit shares striking similarities with the mammalian ventral tegmental area (VTA) dopamine network that suppresses sleep and amplifies courtship during sexual arousal^76–78^. Thus, the neural logic for prioritizing mating over sleep is deeply conserved.

## METHODS

### Fly lines and rearing

All *Drosophila melanogaster* strains were reared on standard cornmeal media at 25°C under 12h:12h light-dark (LD) conditions. Additional dietary supplements were provided for specific species: rotten banana for *Zaprionus indianus*, 7-dehydrocholesterol for *Drosophila pachea*, and noni juice for *Drosophila sechellia*. Species like *D. busckii*, *D. hydei*, *D. mercatorum*, *D. virilis*, *D. americana*, *Z. indianus*, and *D. funebris* were reared at 18°C in 12:12 LD conditions and transferred to 25°C at least 24 h prior to behavioral experiments. All the genetic crosses, except when tub-Gal80^TS^ or TrpA1 was present, were reared at 25°C. For genetic lines with tub-Gal80^TS^ and TrpA1, the flies were reared at 18°C. To inhibit synaptic transmission, we used UAS-TNT recombined with tsh-Gal80, except when targeting VNC neurons. For the majority of the experiments, around 4-5 days old flies were selected from mixed-sex groups of similar densities where the flies were invariably mated within 2-3 days post-eclosion. For species such as *D. virilis*, *D. americana, D. sechellia,* and *D. pachea,* which exhibit delayed sexual maturation, flies around 10-12 days old were used for experiments. The fly lines used in this study are listed in Supplementary Table 2.

### Seminatural assay conditions

Locomotor activity under seminatural conditions was monitored in a garden within the INRAE Versailles campus (48.8021° N, 2.0873° E) situated well away from human disturbance. For the behavioral assays, flies were placed in Trikinetics activity monitors shielded from direct sunlight and rainwater by positioning them in an area under shade. The setup consisted of an enclosure (35 cm × 24 cm × 33 cm) covered at the top, while allowing sufficient air circulation throughout the chamber (Extended Data Fig. 1a). The monitors were connected to a computer located inside the laboratory using long cables. Daily profiles of light, temperature, and humidity were recorded using DEnM devices (Trikinetics, USA). To acclimate the flies to the outside environment, they were housed in standard food vials placed within the same setup for at least one day before starting the seminatural experiments. Various sets of experiments were conducted across seasons from mid-April to mid-October in 2022, 2023, 2024, and 2025.

### Locomotor activity monitoring

Locomotor activity of the adult flies was monitored under seminatural and laboratory conditions using Trikinetics DAM5H (for individual flies and pairs) and Trikinetics LAM25H-3 (for fly groups). Single-beam actigraphic monitors (DAM2) were not consistently able to detect sociosexually driven peaks in locomotor activity. For DAM5H experiments, flies were placed in Pyrex glass tubes (5 mm × 80 mm), with one end filled with food containing 6% sucrose and 2% agar, and the other end sealed with cotton. For LAM25H-3 experiments, flies were housed in polycarbonate tubes (22 mm × 95 mm), one end containing food (6% sucrose and 2% agar), and the other end was sealed with foam plugs. Laboratory experiments were conducted in incubators maintained at 25°C, 60-70% RH under 12h:12h light-dark (LD) cycles (lights on at 10:00 a.m. and off at 10:00 p.m.) with a light intensity of approximately 500-1000 lux. Activity data were binned at 1 min or 30 min intervals and extracted using DAMFileScan114 software (TriKinetics, USA). Activity data were analyzed and visualized using custom Excel scripts and FaasX 1.22 software (available at https://neuropsi.cnrs.fr/wp-content/uploads/2022/09/Faas_Kit.zip). To calculate the N-index and D-index, we adapted the ratio-based scoring described for morning/evening anticipation measurement by Harrisingh et al., 2007^79^. For fly group experiments conducted in LAM25H-3 under both laboratory and seminatural conditions, the N-index was defined as the ratio of average activity/min during the scotophase to average activity/min over 24 h. For our subsequent experiments with fly pairs, we noted that in the opposite-sex pairs, the night peak invariably occurred around the middle of the night. Therefore, for fly pairs, the N-index was calculated as the ratio of the average 1 min bin activity during ZT 14–22 to the same over the entire 24 h day. Similarly, in different Drosophila species pairs, the daytime-specific increase of activity, if present, was restricted within ZT0-8, allowing us to calculate the D-index as the ratio of average 1 min bin activity during ZT0–8 to the same over the entire 24 h. Rhythmicity analysis in FaasX involved chi-square periodogram and autocorrelation methods with the following criteria: for chi-square, filter ON, minimum power ≥ 20, and minimum width ≥ 1.5 h; for autocorrelation, filter ON, and max lag: 144^80^. Nighttime sleep duration in flies was quantified using the MATLAB-based SCAMP program, with sleep defined as a minimum of 5 min of continuous inactivity^81^. All behavioral experiments were reproduced 2-3 times with consistent results. For all analyses, behavioral data from the experimental days 2 and 3 were taken into account. For conditional activation of neurons using UAS-TrpA1, behavioral data recorded at 30°C are shown, and for conditional inhibition using UAS-Kir,tub-Gal80^TS^, behavioral data recorded at 25°C following 3 days of incubation at 30°C are shown.

### Video tracking of the fly behavior

For the analysis of fly activity through video recordings, three pairs of male-female *D. melanogaster* Canton S (3–4-day-old) were housed in each of the wells of a 12-well plate containing a layer of food (6% sucrose and 2% agar) at the bottom, 24 h prior to the start of recording. The plate was covered with a thin, perforated plastic sheet to allow air circulation. It was then placed in a walk-in incubator maintained at 25°C, 60– 70% relative humidity, and a 12h:12h light-dark cycle. Recordings were conducted from ZT0 using a Sony Handycam HDR CX740, utilizing its built-in infrared light for the scotophase. The 24 h recordings were manually analyzed at 30 min intervals for 10 s, and the number of active flies was recorded (Extended Data Fig. 2a).

In order to generate a 24 h ethogram of the fly pairs (Extended Data Fig. 2d), the different behaviors manifested by the heterosexual pairs were analyzed via video recording. Flies were placed in 5 mm DAM5H tubes containing behavioral food within the same arena described above. Video recordings were analyzed manually at 5 min intervals to document behaviors such as interaction, feeding, grooming, micro-movements, ambulation, and resting.

### Remating rhythm

The protocol for the remating assay was modified after Krupp et al., 2013^82^. 3-4-day-old mated *D. melanogaster* Canton S males and females were separated into same-sex groups for 24 h prior to the experiment. Two pairs of male and female flies were placed in each well of a 24-well plate containing a layer of cornmeal agar food at the bottom. The plate was then covered with a thin plastic sheet perforated with tiny holes to allow ventilation and placed in a walk-in incubator maintained at 25 °C, 60–70% relative humidity, and a 12 h:12 h light–dark cycle. On the following day, fly pairs were continuously recorded for 24 h starting from ZT12 using a Sony Handycam HDR-CX740, with its infrared mode enabled during the scotophase. Recordings were manually analyzed at 2 min intervals to score remating events.

### Courtship assays

To analyze the courtship index, mated males and females of different Drosophila species were individually introduced into DAM5H tubes containing food (6% sucrose and 2% agar) at one end. Flies were transferred using gentle suction to avoid anesthesia, and were acclimated for at least 12 h prior to the start of the experiment. Only for the experiments to check the acute courtship index of TH-F2>TNT, tsh-Gal80 flies (Extended Data Fig. 7i), virgin males and females were used. The tubes were placed in a walk-in incubator maintained at 25°C, 60–70% relative humidity, and a 12h:12h light-dark cycle. Immediately before the onset of the recording, males and females were introduced into the same DAM5H tube. Around 15 min of video recordings were conducted at defined time points using a Sony Handycam HDR CX740, utilizing its built-in infrared light for the recordings in the dark. The recordings were manually analyzed at 10 s intervals for a total of 10 min, noting the number of pairs engaged in species-specific tell-tale signatures of courtship behavior. The courtship index was calculated using the following formula: Courtship Index (CI)=Total time engaged in courtship/Total observation time^83^.

### Close-proximity assay

The protocol of the close-proximity assay was adapted from Hamasaka et al., 2010^19^. Transparent 35 mm petri dishes were filled with food containing 6% sucrose and 2% agar, leaving a 3 mm gap between the food surface and the top edge of the dish. Around 3–4 days old, Canton S male-female pairs were introduced into each petri dish around dusk. The dishes were arranged on a platform designed to diffuse uniform infrared light (850 nm), positioned under a USB camera (USB4K02AF-V100) for imaging. Each dish was covered with a thin transparent glass plate, leaving a 0.5 mm gap to ensure adequate ventilation. The entire experimental setup was placed in an incubator maintained at 25°C, 60-70% relative humidity, and a 12h:12h light-dark cycle. The image acquisition system consisted of a Raspberry Pi 4B nano-computer connected to the USB camera. Image acquisition started at ZT12 and continued for 24 h, during which high-resolution images (1920px × 1080px) were captured every 10 s. Captured images were analyzed using a custom Python script with OpenCV libraries. The analysis began with identifying the outlines of the Petri dishes in the first image to generate masks, in which automatic thresholding and binarization were applied to each frame to identify the positions of individual flies. For each frame, the distance (d) between the two flies in a pair was calculated. A score of 0 was assigned when d>5 mm and a score of 1 when d≤5 based on Fujii et al., 2007^18^. For each pair, proximity scores were summed over 2 h intervals and expressed as a percentage of the maximum possible interactions (120 min x 6 images/min = 720 interactions). However, it’s worth noting that a singular proximity score may not be enough to denote sociosexual interactions^84^.

### Cuticular hydrocarbon analysis using GC-MS

The protocol for cuticular hydrocarbon analysis was modified after Kuo et al., 2012^85^. The qualitative and quantitative analysis of the cuticular hydrocarbons (CHC) of *D. melanogaster* strains was achieved using a solvent extraction method in gas chromatography-mass spectrometry (GC-MS). Female flies, aged 6–7 days, were separated the night before extraction. 60 flies were placed in a 2 mL glass vial, and 1 mL of *n*-hexane was added to it. The vial was vortexed thoroughly for 20-30 s and left to stand for 30 minutes. The extract was then recovered and transferred to a chromatography vial, and 10 μL of an *n*-eicosane solution (2 mg/L) was added as an internal standard. A 1 μL aliquot of the sample was injected into an Agilent 8890 GC system coupled to an Agilent 5977B single quadrupole mass spectrometer. The injector was operating in splitless mode at 250°C. The chromatographic separation was achieved using an HP-5 MS capillary column (30 m × 0.25 mm × 0.25 μm; Agilent Technologies) with helium as the carrier gas at a constant flow rate of 1.5 mL/min. The oven was heated from 50°C to 300°C with a ramp of 9°C/min. Ionization was performed in electron impact mode (70 eV), and a full scan mode was used with an m/z range of 35–420. The source and quadrupole temperatures were maintained at 230°C and 150°C, respectively. GC-MS data were processed using Agilent MassHunter software, and the compounds were identified by comparing their mass spectra using the NIST Tandem Mass Spectral Library (version 2.4). Retention indices were calculated using a saturated n-alkane (C20–C30) standard solution. A quantitative analysis of the identified compounds of interest was done by dividing their integrated peak areas by that of the internal standard. The extraction and injection were repeated three times, and the mean relative abundance of the compounds of interest was subsequently computed.

### Feeding the flies with the agonist and antagonist of dopamine signaling

A 1 mg/ml solution of the 3-Iodo-L-tyrosine (3IY) (Sigma-Aldrich; stored at –20 °C) was prepared by dissolving the compound in distilled water, and the solution was supplemented with 6% sucrose. Prior to the administration of the 3IY solution, the flies were wet-starved overnight, during which they were maintained without food but provided with access to water. For 3IY administration, 2 ml of the 3IY solution was dispensed onto a Kimwipe paper placed inside an empty Drosophila vial. The wet-starved flies were then transferred to these vials and allowed to feed on the solution for 48 h before running behavioral tests.

For L-3,4-dihydroxyphenylalanine (L-DOPA) administration, wet-starved flies were fed using the same procedure with a solution containing L-DOPA (2 mg/ml, Sigma-Aldrich) and Carbidopa (0.05 mg/ml, Sigma-Aldrich) for the initial 48 h before experiments. During L-DOPA feeding, the fly vials were kept well protected from bright light to prevent compound degradation.

For both 3IY and L-DOPA treatments, the flies were also maintained on the same respective diets throughout the duration of the behavioral experiments.

### Optogenetic studies

Optogenetic experiments were conducted using an ethoscope-based tracking system, as previously described by Geissmann et al., 2017^86^. Experimental flies were fed a solution of 800 μM all-trans-retinal (ATR; Sigma-Aldrich) mixed with standard agar food for three consecutive days in complete darkness. After this period, individual flies from both ATR-fed and unfed (control) groups were transferred to custom-made rectangular arenas (6 cm × 5.5 cm × 0.3 cm), sealed with transparent, anti-reflective glass covers. The arenas were then mounted inside ethoscopes for independent tracking of each fly.

Ethoscopes were placed in an incubator maintained at 25°C and illuminated with ambient infrared light (850 nm) to enable tracking in darkness. Each fly was recorded across three distinct phases. In Phase 1 (pre-stimulation), flies were tracked in complete darkness for 1 h. In Phase 2 (stimulation), a red spotlight (600 nm) positioned at the center of each arena was activated to optogenetically stimulate the flies while tracking continued for 1 h. In Phase 3 (post-stimulation), the red light was turned off, and flies were tracked in darkness for an additional h. Behavioral data were analyzed using a custom-written Python script. Fly position data from each phase were extracted from ethoscope-generated database files (.db), and two-dimensional heatmaps were generated to visualize spatial occupancy patterns. Light-zone preference was quantified as the difference between the percentage of time spent in the red-light spot and that in a proximal fictitious spot of equal size.

### CHC preference assay

The cuticular hydrocarbon (CHC) cocktail was extracted from 60 *D. melanogaster* Canton S females using the solvent extraction method described above. 1 ml of the female CHC extract was evenly spread over the surface (∼7 cm²) of solidified behavioral food (6% sucrose, 2% agar) and allowed to air dry until complete evaporation of the solvent hexane. The CHC-coated food was then inserted into one end of individual DAM tubes (5 mm × 65 mm) such that the CHC-treated surface faced inward. Following this, 3-4-day-old virgin Canton S males were singly inserted into each tube. The opposite end of each tube was sealed with untreated behavioral food (without CHCs). A tiny hole was present in the middle of each tube to allow air circulation. The tubes were placed in a TriKinetics DAM5H activity monitors housed in an incubator maintained at 25 °C, 60–70% relative humidity, and a 12 h:12 h L:D cycle, starting around ZT12. The individual fly’s position was recorded using the Dwell Quad module of the DAMSystem3 software, which divides each tube into four equal quadrants (D1–D4). Positional preference toward the CHC-coated food was quantified using the preference index formula: Preference Index = (Dwell in CHC quadrant – Dwell in opposite end quadrant) /Total dwell.

### Connectome mining

Connectome searches were conducted using the neuPrint platform (https://neuprint.janelia.org/). Pair-wise connectivity of PPM2 and pCd1 neurons was mined using both the female (hemibrain:v1.2.1) and male (male-cns:v0.9) connectome datasets. However, direct synaptic connectivity was detected only in the male CNS. Within the male pCd1 population, the SMP721m neuron was identified as a direct synaptic output of PPM1201 (PPM2). This connection was further validated using neuPrint’s ‘shortest path’ query search with body IDs 12644 (PPM1201) and 81351 (SMP721m).

For anatomical validation and visualization, 3D reconstructions of PPM2 and pCd1 neurons were generated using the Neuroglancer and Skeleton viewer tools available in the neuPrint interface. Neuron skeletons and spatial reconstructions were then downloaded from the male-cns:v0.9 dataset for publication.

### GCaMP imaging

GCaMP imaging protocol was adapted from Chatterjee et al., 2025^53^. Adult *D. melanogaster* brains were dissected in ice-cold HL3 (hemolymph-like solution). Whole-brain explants were placed in a custom-made imaging chamber pre-treated with Poly-D-lysine and immersed in 700 µL of HL3 solution. Calcium imaging was performed using a two-photon microscope (Karthala AODscope system) with GCaMP6s fluorescence excited at 920 nm. P2X₂ channel activation was achieved by applying ATP (Sigma-Aldrich) to the bath solution via a microsyringe, resulting in a final ATP concentration of 5 mM. Baseline calcium activity was recorded for 1 min prior to ATP application, followed by 4 min of imaging after ATP addition. Image stacks (xyzt) were acquired and converted to maximum intensity projections to generate static images. For fluorescence quantification, the mean fluorescence intensity of all pixels within a defined region of interest (ROI) was measured for each time point, and background fluorescence was subtracted using a region devoid of signal within the same brain. The resulting background-corrected fluorescence at each time point was defined as F_b_. Changes in fluorescence were computed as %ΔF/F₀ = ((F_b_−F₀)/F₀) ×100, where F₀ represents the mean background-corrected baseline fluorescence during the 1 min period preceding ATP stimulation. Image processing and quantification were performed using Fiji (ImageJ). The maximum change in GCaMP6s fluorescence (Max ΔF/F₀) was determined as the maximum percentage change observed during each recording. For each genotype, maximum values from individual samples were averaged to obtain the mean maximum fluorescence change from baseline.

### Immunostaining

The immunolabelings were performed using whole-mounted adult fly brains as described by Chatterjee et al., 2025^53^. Briefly, brains were dissected in cold PBS, fixed in 4% PFA for 1h at room temperature, washed 6x in 10 min steps with PBST (PBS containing 0.3% TritonX), and permeabilized with PBS containing 1% TritonX before undergoing overnight blocking in 1% BSA solution. Primary antibodies – anti-GFP (chicken; 1:1000, ThermoFisher, Cat# A10262), anti-HA (rat; 1:200, Roche, Cat# 11867423001), anti-TH (rabbit; 1:200, ThermoFisher, Cat# OPA1-04050), anti-nc82 (mouse; 1:200, DSHB, Cat# nc82) were diluted in PBST containing 0.1% BSA and used for 2-3 nights at 4°C. After washing, brains were treated with secondary antibodies for 2-3 nights at 4°C, washed with PBS containing thimerosal, and mounted in Vectashield. Secondary antibodies (at 1:1000 dilution) used were: goat anti-chicken IgY Alexa 488 (Thermo Fisher, Cat# A-11039), goat anti-rat IgG FP 546 (Interchim, Cat# FP-GARAT546), goat anti-rabbit IgG FP 568 (Interchim, Cat# FP-GARB568), and goat anti-mouse IgG Alexa 647 (Thermo Fisher Scientific, Cat# A-21235). Fluorescence signals were acquired with Zeiss Zen2 software from an AxioImager Z1 semi-confocal microscope equipped with an AxioCam MRm digital camera and an ApoTome2 module set with an automatically adjustable grid, which provided structured illumination. Depending on the experiment, 25x or 40x objectives were used. Fiji software was used to quantify fluorescence intensity from digital images. For all quantifications from soma (CaLexA experiments), we utilized the formula: I = 100x(S-B)/B to calculate the fluorescence percentage above background (S = mean intensity inside the cell, B = the mean intensity of the region adjacent to the positive cell). For quantifications from neuropils, integrated densities over a defined area were measured.

### Statistics and Reproducibility

All statistical analyses were performed using Prism (version 10, GraphPad Software, USA) and R (version 4.3.0; R Core Team). All measurements were taken from distinct biological samples, and no individual sample was measured repeatedly.

For comparing two sample means, a two-tailed unpaired t-test with Welch’s correction, and for multiple sample means, one-way ANOVA followed by a post hoc test was performed. Differences in the % rhythmicity and phase vectors between groups were assessed using a Z-test for population proportions and Watson’s U2 test, respectively. Correlations between proximity and velocity (Extended Data Fig. 2g) were assessed using Pearson’s product–moment correlation test using Pearson’s correlation coefficient. Effect sizes for mean comparisons (Fig. 2a) were quantified using Cohen’s *d*, calculated via custom R scripts.

Cluster analyses were performed using the ClustVis web tool (https://biit.cs.ut.ee/clustvis/). Non-stationary Hidden Markov Model (HMM) analysis, principal component analysis (PCA), K-means clustering, and canonical correlation analysis (CCA) were implemented using custom R scripts. The phylogenetic tree of 21 Drosophila species was reconstructed based on data from Suvorov et al., 2022^87^ using custom R code (Extended Data Fig. 3c). Phylogenetic signal in the temporal niche patterning of the species was evaluated using Pagel’s λ test. In all cases, statistical significance was defined as p< 0.05.

## Supporting information

statistics table

## DATA AVAILABILITY

Numerical source data underlying the main and Extended Data figures have been deposited in the Figshare repository under DOI: 10.6084/m9.figshare.33111071^88^. Due to their large file sizes, raw behavioral activity log files and microscopy images were not deposited in the public repository, but are available from the corresponding author upon reasonable request. Source data are also provided with this paper.

## CODE AVAILABILITY

Custom codes used for data analysis and statistical processing are included in the Source Data package provided with this paper and have also been deposited in the Figshare repository under DOI: 10.6084/m9.figshare.33111071^88^.

## ACKNOWLEDGEMENTS

We thank G. Thijs, S. Hirschmann, A. Païta, H. El Hajj, S. Aryal, Z. Chak, and P. Couzi for their help with various experiments; A. Yassin, C. Helfrich-Förster, P. Deppisch, T. Auer, R. Benton, M. Noll, J.C. Billeter, D. Vasiliauskas, T. Preat, F. Martin, M. Vasconcelos and S. Vasu for sharing fly stocks with us; C. Bustos, C. Fabre and P. Lucas for their valuable discussions and help with data analyses; P.E. Hardin, C. P. Kyriacou, and J.C. Billeter for their insightful comments on an earlier version of the manuscript. We dedicate this work to P. K. Chattopadhay.

## AUTHOR CONTRIBUTIONS

A.C. designed the project and secured funding; S.G. and A.C. designed the experiments. S.G., Z.P., C.S., T.C., J.M., and V.R. performed behavioral experiments; S.G. and Z.P. did the immunostainings and GCaMP imaging; A.P. and S.G. did the GC-MS experiments. S.G. and A.C. analyzed the data with the help of others. F.R. co-supervised Z.P. and T.C. with A.C. A.C. and S.G. wrote the manuscript with input from other authors.

---

**Supplementary Figure 1.**
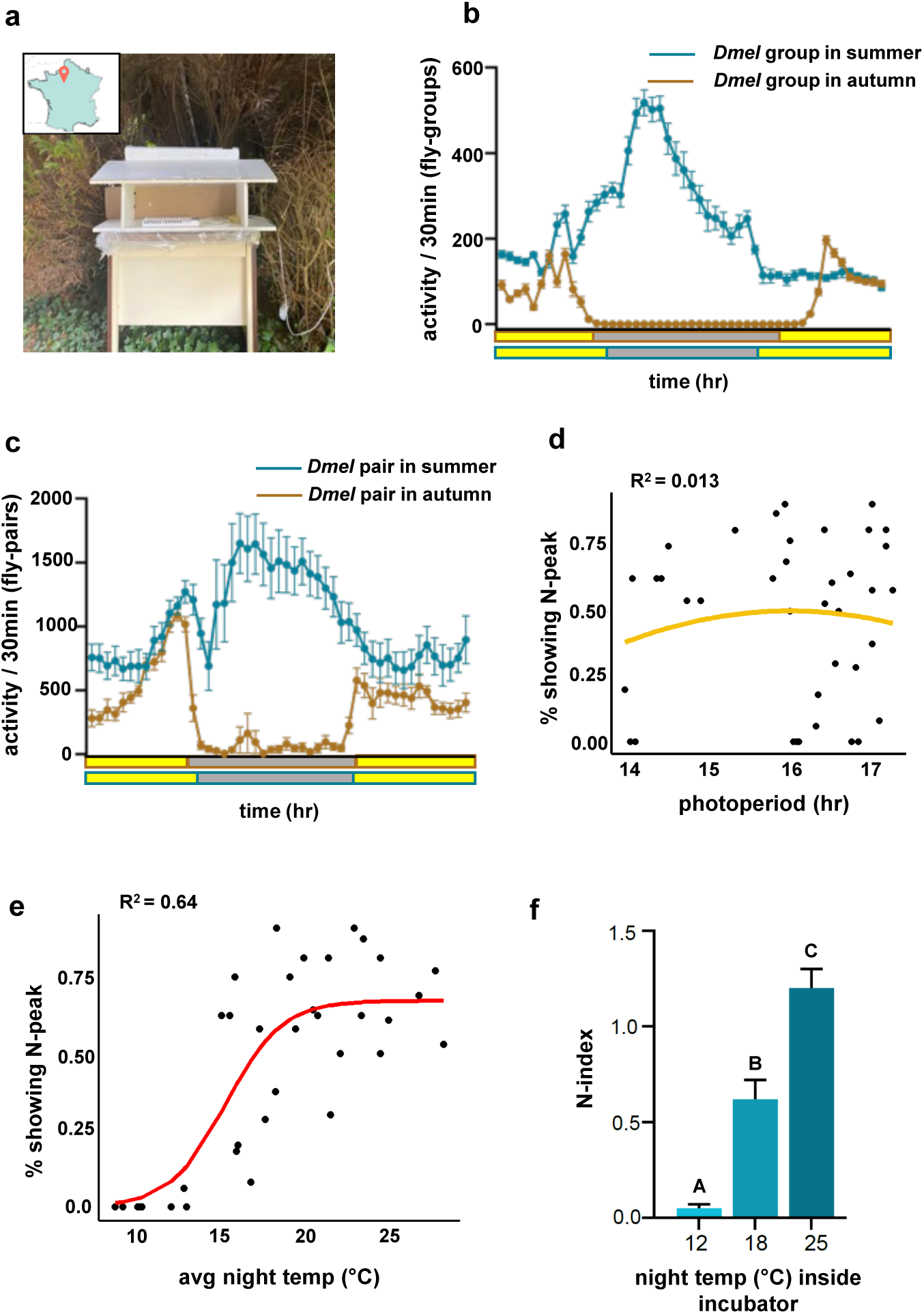

**Supplementary Figure 2.**
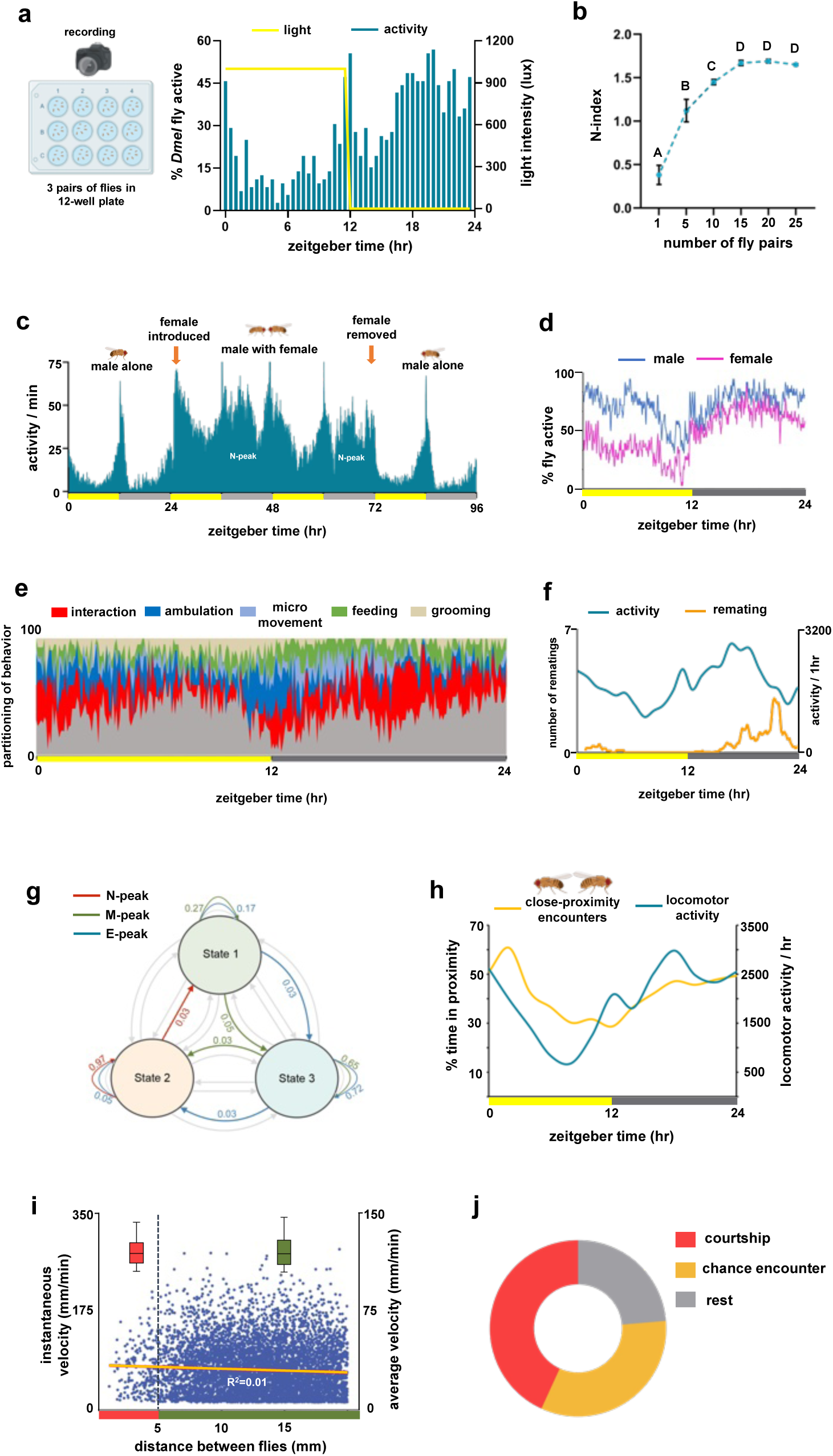

**Supplementary Figure 3.**
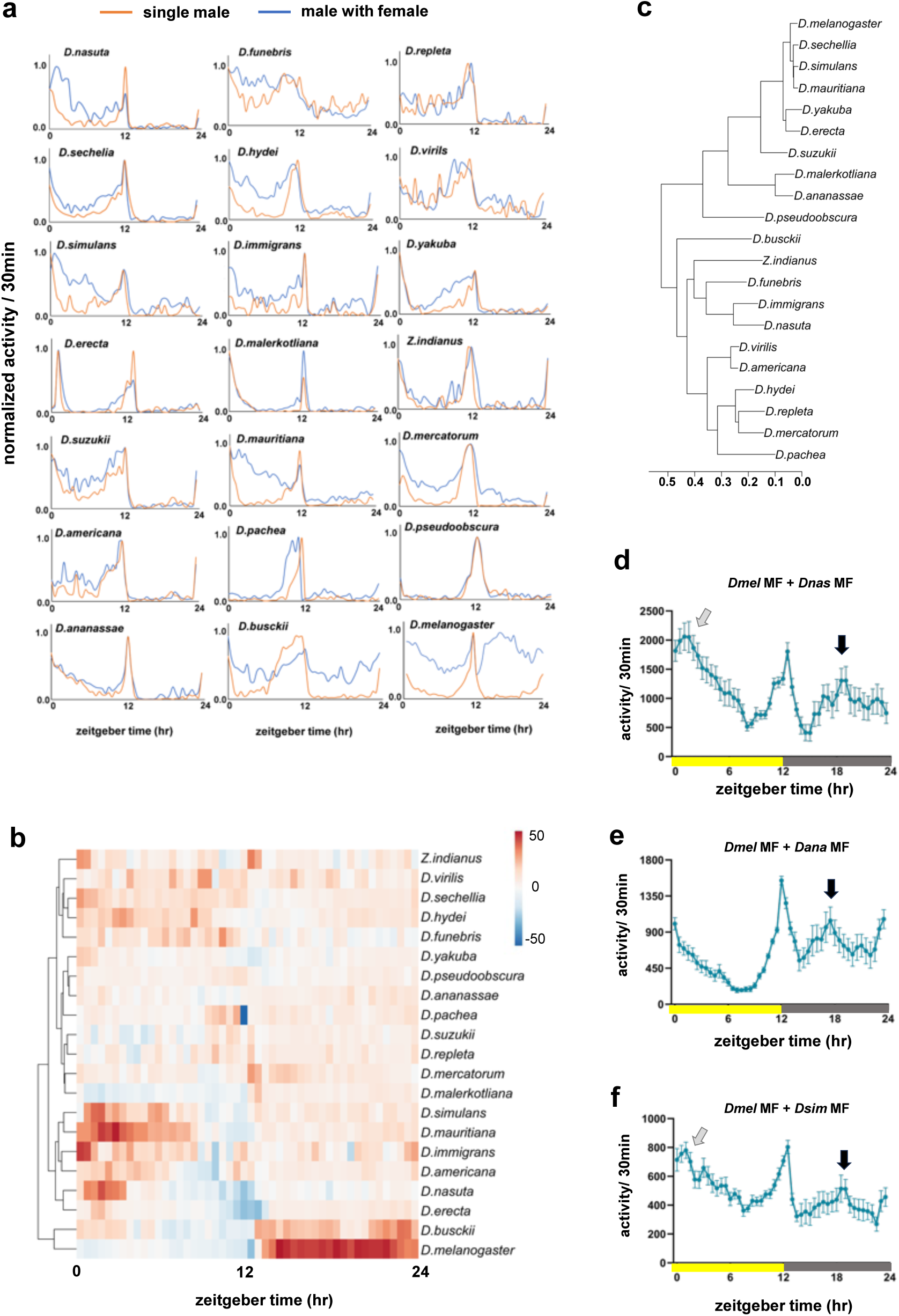

**Supplementary Figure 4.**
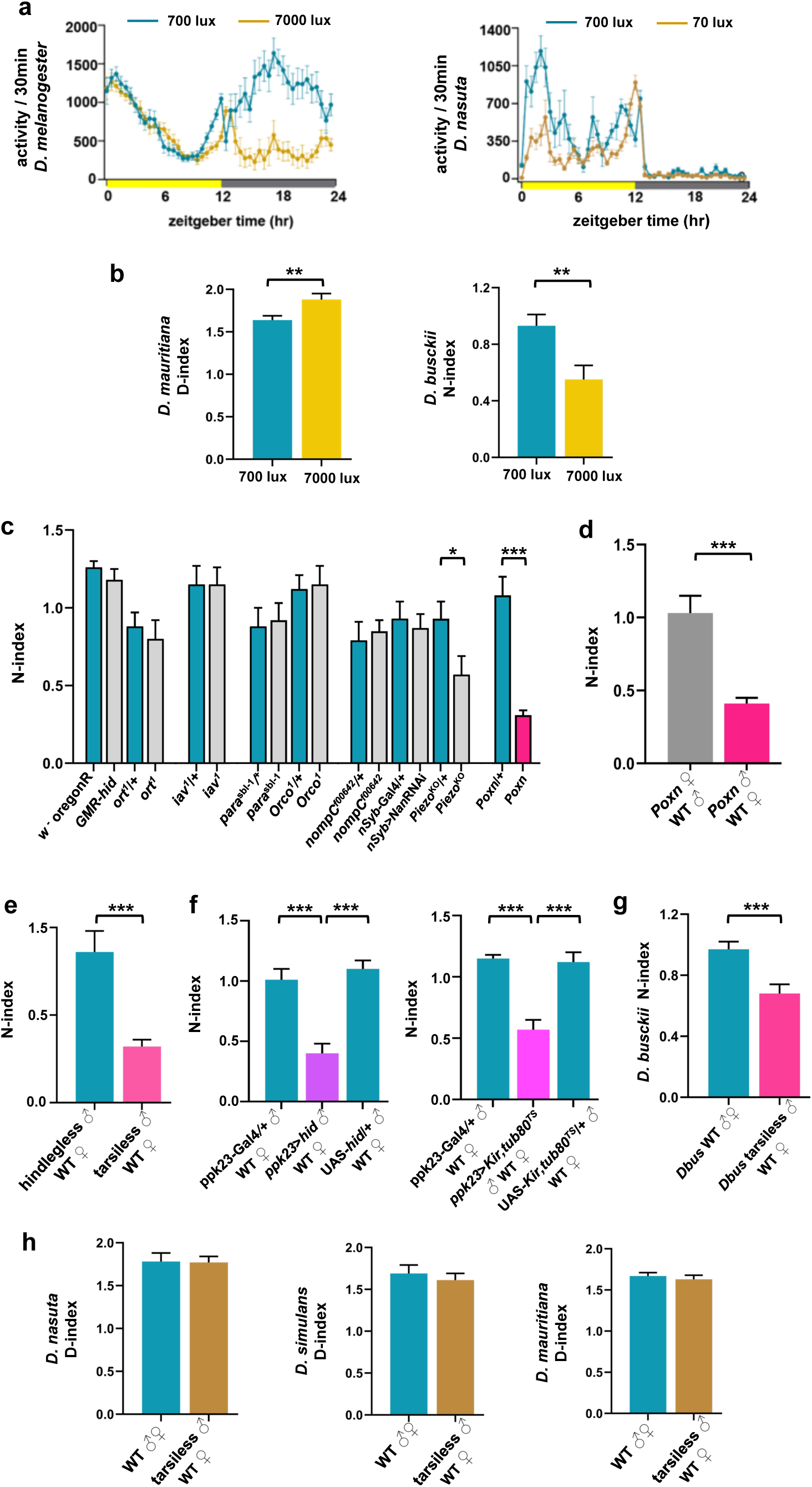

**Supplementary Figure 5.**
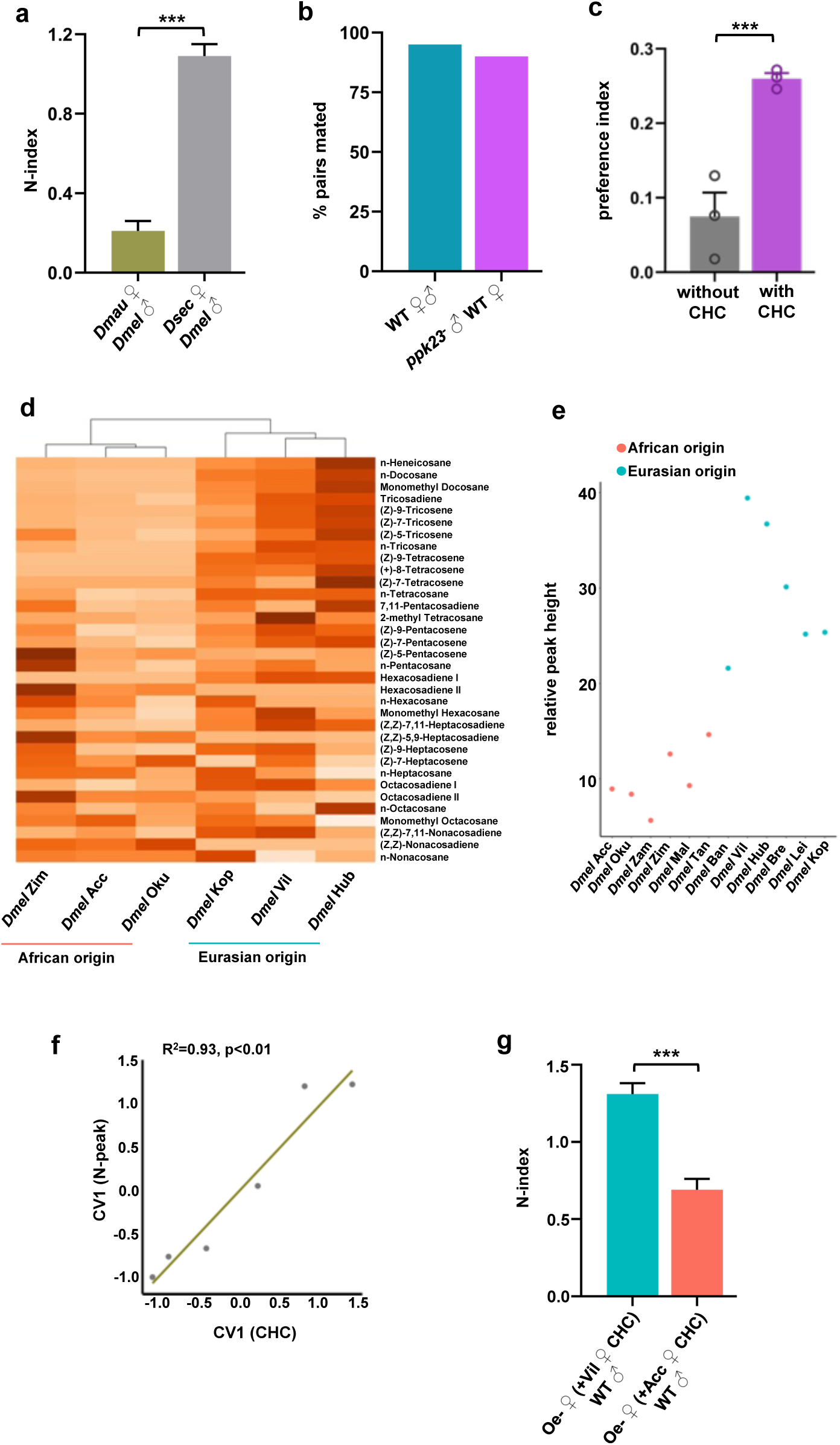

**Supplementary Figure 6.**
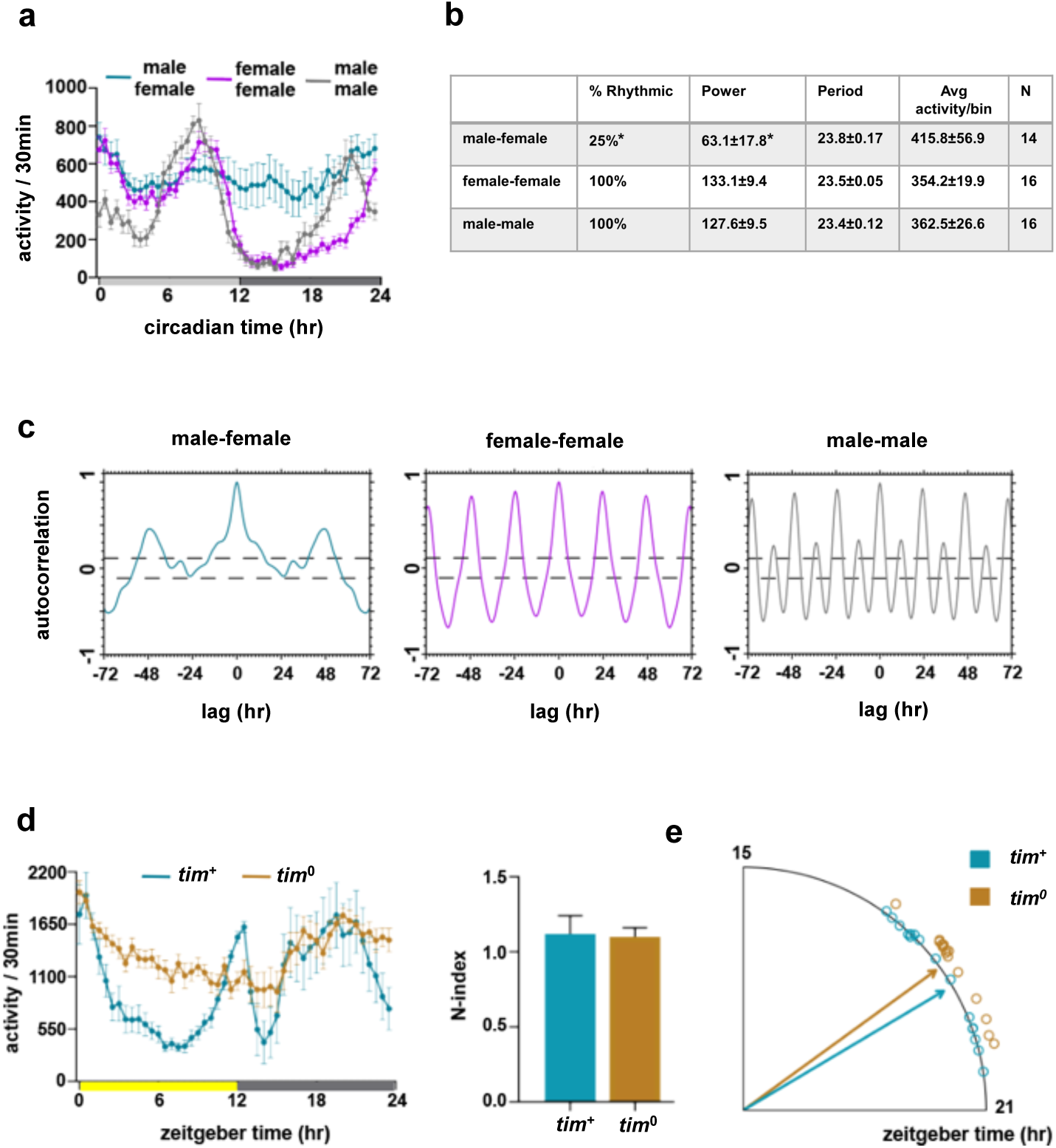

**Supplementary Figure 7.**
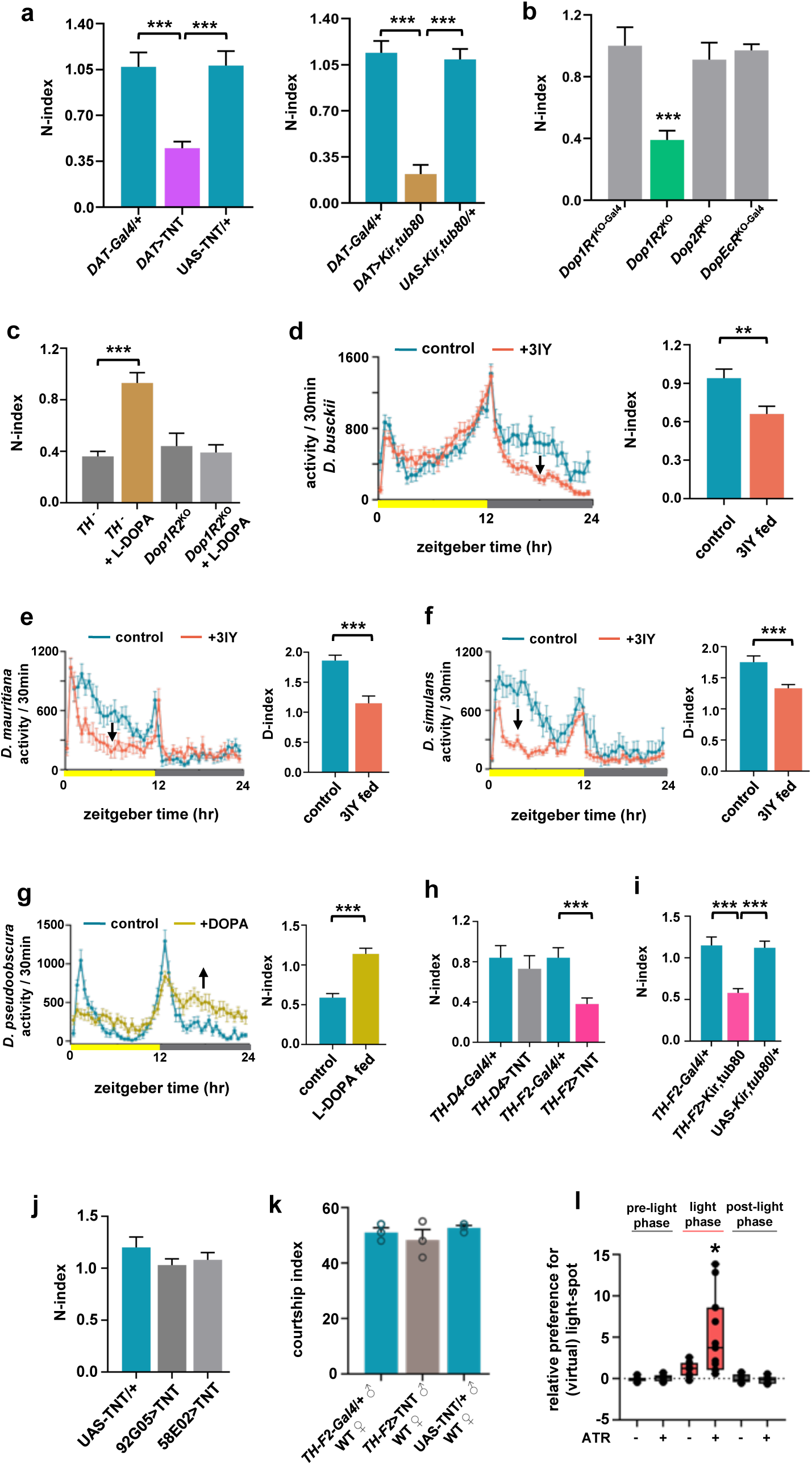

**Supplementary Figure 8.**
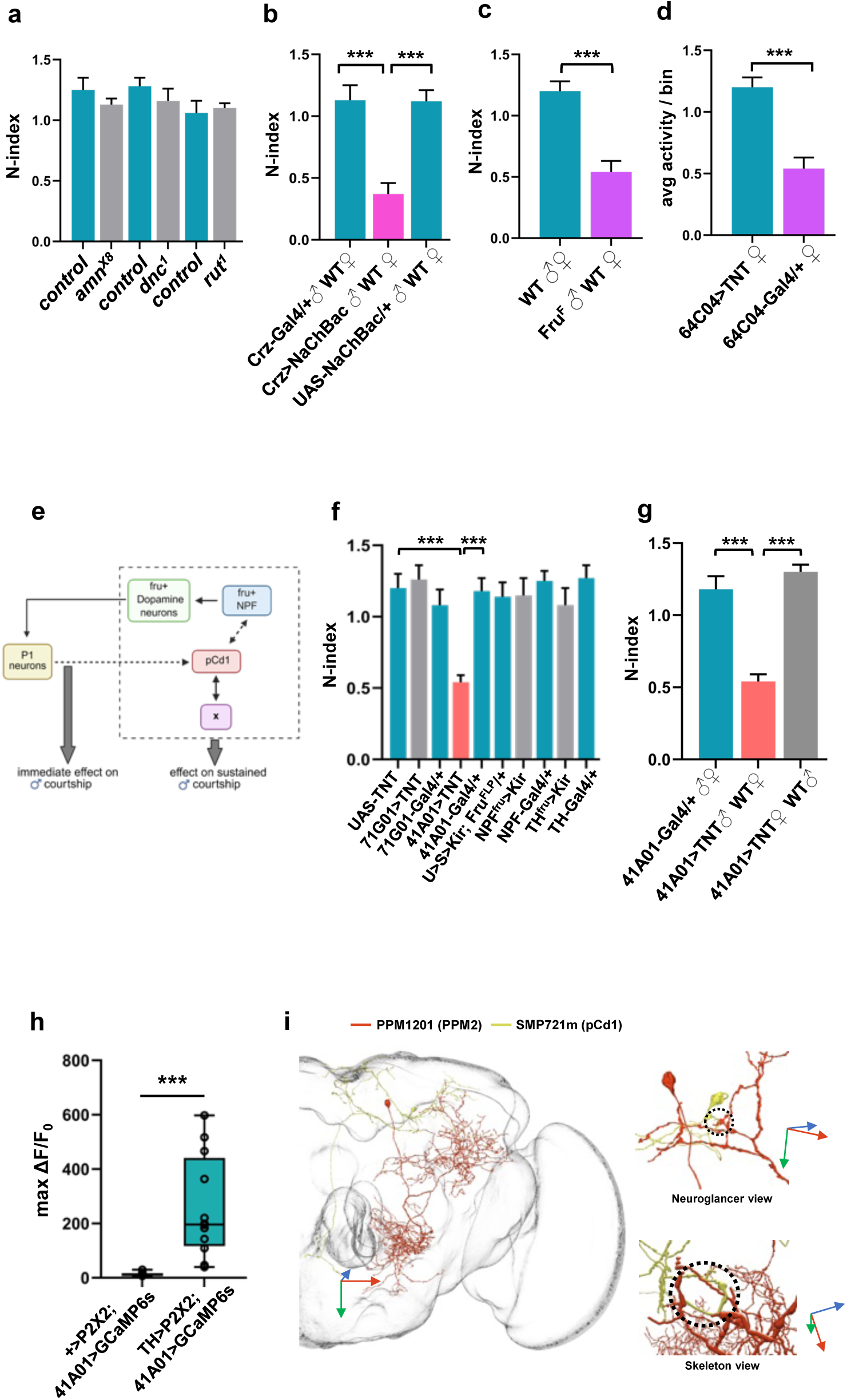

**Supplementary Figure 9.**
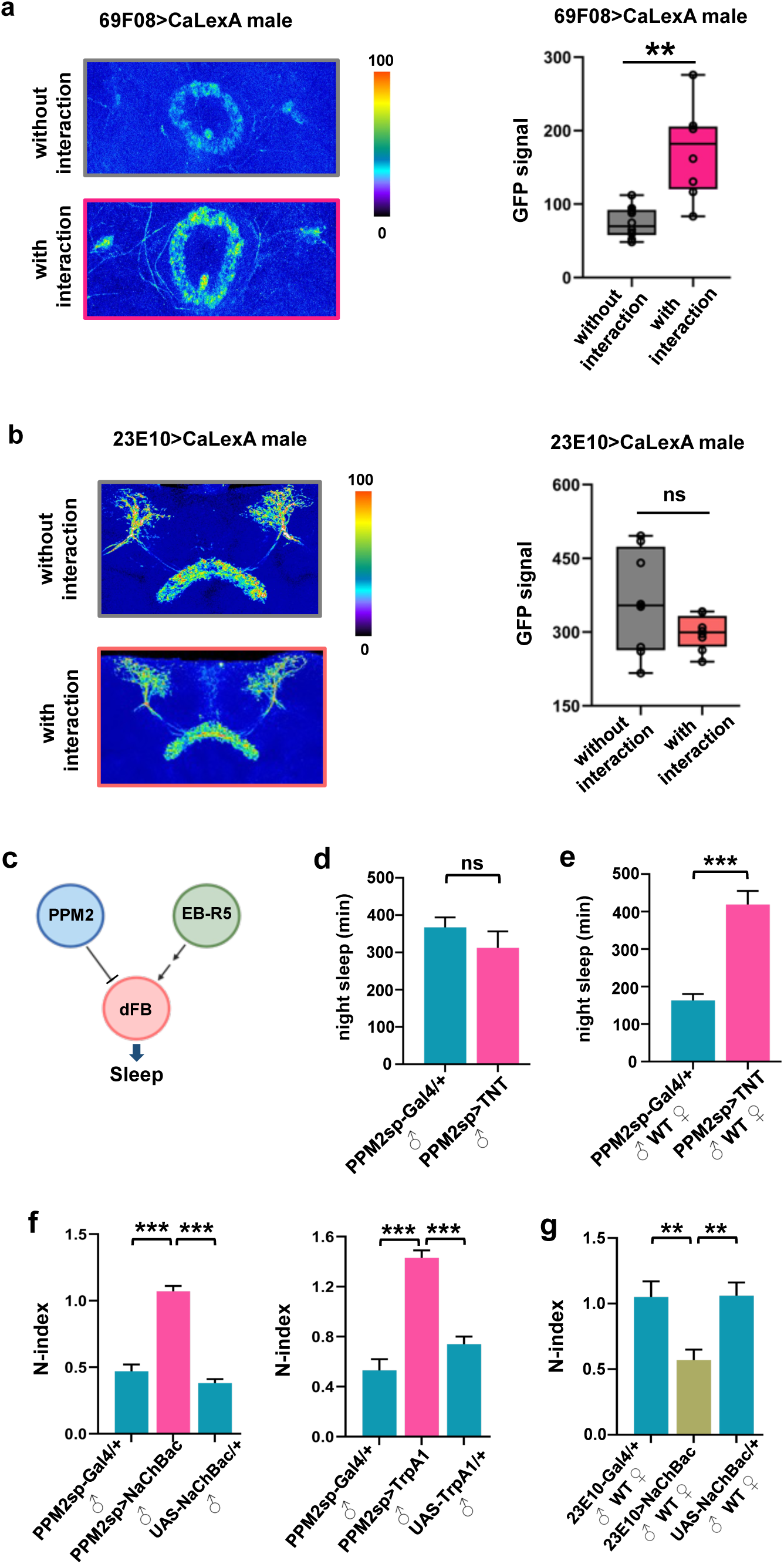

