## supplementary information for "Mating imperatives drive plasticity of the daily temporal niche in fruit flies"

### SUPPLEMENTARY TABLES:

**Supplementary Table 1**: Detailed statistics for main and extended data figures.

**Supplementary Table 2**: List of fly species, strains, and transgenic lines used.

**Supplementary Table 3:** Details on genetic manipulations done in the manuscript.

**Supplementary Table 4:** Behavioral features associated with HMM-inferred states.

### SUPPLEMENTARY VIDEOS:

**Supplementary Video 1**: Aggregation of mixed-sex *D. melanogaster* flies on a food patch (~8 cm²) at an approximate density of 11 flies per cm² (N=100 flies).

**Supplementary Video 2:** Wild-type *Dmel* male-female pair showing courtship behaviours and copulation in our assay system.

**Supplementary Table 2:** Genetic reagents and *Drosophila* species used in this study.

| **Genotype** | **Source** |
|---|---|
| **Wild type** | |
| *D.mel* Canton S | BDSC#64349 |
| *D.mel* $w^-$*; OregonR* | Jean-Christophe Billeter, University of Groningen |
| *D.mel* $w^{1118}$ | BDSC#3605 |
| *D.mel Viltain* | EGCE, IDEEV |
| *D.mel Hub11* | Peter Deppisch, University of Würzburg |
| *D.mel Bret1* | Daniel Vasiliauskas, NeuroPSI |
| *D.mel Kop3* | Peter Deppisch, University of Würzburg |
| *D.mel Lei38* | Peter Deppisch, University of Würzburg |
| *D.mel Accra1* | Peter Deppisch, University of Würzburg |
| *D.mel Oku* | EGCE, IDEEV |
| *D.mel Zambia* | Peter Deppisch, University of Würzburg |
| *D.mel Zimbabwe S-29* | BDSC#60741 |
| *D.mel Tanzania* | Peter Deppisch, University of Würzburg |
| *D.mel Malawi 63* | BDSC#32046, BDSC#30864 |
| *D.mel Bangalore* | Sheeba Vasu, JNCASR |
| *D.yakuba* | EGCE, IDEEV |

| | |
|---|---|
| *D.mauritiana 90* | Richard Benton, University of Lausanne |
| *D.malerkotliana* | EGCE, IDEEV |
| *D.americana texana* | Peter Deppisch, University of Würzburg |
| *D.simulans* | EGCE, IDEEV |
| *D.immigrans* | EGCE, IDEEV |
| *D.virilis* | Peter Deppisch, University of Würzburg |
| *D.pachea* | Michael Lang, EGCE |
| *Z.indianus* | Sheeba Vasu, JNCASR |
| *D.suzukii* | Jean-Luc Gatti, ISA |
| *D.sechellia* | Richard Benton, University of Lausanne |
| *D.erecta C3* | EGCE, IDEEV |
| *D.mercatorum* | Charlotte Förster, University of Würzburg |
| *D.nasuta* | Sheeba Vasu, JNCASR |
| *D.funebris* | Peter Deppisch, University of Würzburg |
| *D.hydei* | Peter Deppisch, University of Würzburg |
| *D.pseudoobscura* | EGCE, IDEEV |
| *D.ananassae* | EGCE, IDEEV |
| *D.repleta K* | Peter Deppisch, University of Würzburg |
| *D.busckii* | Peter Deppisch, University of Würzburg |
| **Mutant** | |
| GMR-hid | François Rouyer, NeuroPSI |
| ort$^{1}$ | François Rouyer, NeuroPSI |
| per$^{0}$ | BDSC#80928 |
| tim$^{0}$ | BDSC#80930 |
| iav | Jean-Christophe Billeter, University of Groningen |
| ppk23$^{-}$ | Jean-Christophe Billeter, University of Groningen |
| para$^{sbl-1}$ | BDSC#60687 |
| Orco$^{1}$ | BDSC#23192 |
| nompC$^{f00642}$ | Jean-Christophe Billeter, University of Groningen |
| piezo$^{KO}$ | BDSC#58770 |
| Poxn-resc$^{\Delta XBs6}$; Poxn$^{\Delta M22-B5}$ | Markus Noll, University of Zurich |
| Ir52c-d$^{-}$ | BDSC#60689, BDSC#60690 |
| Tbh$^{nM18}$ | BDSC#93999 |
| Trh$^{01}$ | BDSC#86147 |
| DTHg$^{FS\pm}$;ple | Serge Birman, ESPCI |
| SPR$^{attp}$ | BDSC#84576 |
| fru$^{4-40}$ | BDSC#66692 |
| fru$^{M}$ | BDSC#66874 |
| fru$^{FLP}$ | BDSC#66870 |
| amn$^{X8}$ | Thomas Preat, ESPCI |
| dnc$^{1}$ | BDSC#6020 |
| rut$^{1}$ | Francisco Martin, Institute Cajal |
| Dop1R1$^{KO-Gal4}$ | BDSC#84714 |
| Dop1R2$^{KO}$ | BDSC#84719 |
| Dop2R$^{KO}$ | BDSC#84720 |
| DopEcR$^{KO-Gal4}$ | BDSC#84717 |
| **Gal4/Split-Gal4/LexA** | |
| Desat1(PromE)-Gal4 | BDSC#65405 |

| | |
|---|---|
| 57C10 (nSyb)-Gal4 | BDSC#39171 |
| *promE-Gal4, tubPGal80ts* | Jean-Christophe Billeter, University of Groningen |
| 55C10AD; ple-DBD$^{THC}$ (PPM2 split) | BDSC#600804, BDSC#95077 |
| 58E02-Gal4 | BDSC#41347 |
| 92G05-Gal4 | BDSC#48416 |
| TH-D4-Gal4 | François Rouyer, NeuroPSI |
| 64C04-Gal4 | BDSC#39296 |
| VT038154AD; dsx$^{DBD}$ (OvAbg split) | Maria Vasconcelos, Champalimaud Foundation |
| 71G01-Gal4 | BDSC#39599 |
| NPF-Gal4 | BDSC#25681 |
| 41A01-Gal4 | BDSC#600669 |
| 69F08-Gal4 | BDSC#39499 |
| 23E10-Gal4 | BDSC#49032 |
| DAT$^{2A}$-Gal4 | BDSC#84622 |
| TH-F2-Gal4 | François Rouyer, NeuroPSI |
| ppk23-Gal4 | BDSC#93026 |
| Crz-Gal4 | BDSC#51977 |
| TH-LexA | François Rouyer, NeuroPSI |
| **UAS/LexAop** | |
| UAS-hid, UAS-Stinger | Jean-Christophe Billeter, University of Groningen |
| UAS-Nanchung RNAi | Caroline Fabre, iEES-Paris |
| UAS-tra2RNAi | BDSC#99704 |
| UAS-TNT | BDSC#28997 |
| UAS-TNT, tsh-Gal80 | Tihana Jovanic, NeuroPSI |
| UAS-NaChBac | BDSC#9469 |
| UAS-CsChrimson | BDSC#55135 |
| UAS-GFP | BDSC#1521 |
| UAS-mCD8GFP | BDSC#5130 |
| UAS-CaLexA | BDSC#86325 |
| UAS-GCaMP6s | BDSC#42746 |
| LexAop-P2X$_2$ | BDSC#76030 |
| UAS-retro-Tango | BDSC#99661 |
| UAS>STOP>Kir2.1 | BDSC#67686 |
| UAS-Dop1R2.gRNA | BDSC#97792 |
| UAS-Cas9.P2 | BDSC#58986 |
| UAS-TrpA1 | BDSC#26263 |
| UAS-Kir2.1, tub-Gal80$^{TS}$ | Tihana Jovanic, NeuroPSI |

**Supplementary Table 3:** Details on genetic manipulations done in the manuscript.

| **Figure number** | **Genotype** | **Explanation** |
|---|---|---|
| Fig. 2e | PromE>traRNAi | Pheromone-producing cells were masculinized in *Dmel*. |
| Fig. 3h | 55C10-AD; ple-DBD$^{TH-C}$ (PPM2-sp) | Genetic line to target a small subset of dopamine neurons (PPM2) in the *Dmel* brain. |

| | | |
|---|---|---|
| Fig. 3h | 55C10-AD; ple-DBD$^{TH-C}$>GFP | PPM2 dopamine neurons were labeled with the fluorescent protein GFP. |
| Fig. 3h | 55C10-AD; ple-DBD$^{TH-C}$>TNT, tsh-Gal80 | Activity of the PPM2 dopamine neurons was silenced in the *Dmel* brain. |
| Fig. 3i | TH-F2>CaLexA | Calcium-dependent reporter was expressed to mark the activation of specific subsets of dopamine neurons (PPM2 & PPM3). |
| Fig. 3l | 55C10∩ple$^{TH-C}$>CsChrimson | PPM2 dopamine neurons in the *Dmel* brain were activated using optogenetics. |
| Fig. 4d | 64C04>TNT | Neurons that regulate *Dmel* females' transient stalling during courtship were silenced. |
| Fig. 4f | VT038154∩dsx (OvAbg-sp) | Genetic line to target neurons that control male abdominal bending (OvAbg). |
| Fig. 4f | VT038154∩dsx>TNT | Abdominal bending-regulating OvAbg neurons were silenced in *Dmel* males. |
| Fig. 4g | 71G01>TNT, tsh-Gal80 | Courtship-promoting P1/pC1 neurons were inhibited in *Dmel* brain. |
| Fig. 4g | TH$^{fru}$>Kir | The activity of a subset of *fruitless*-expressing dopamine neurons involved in courtship was silenced. |
| Fig. 4g | NPF$^{fru}$>Kir | A subset of NPF neurons expressing *fruitless* and involved in courtship was inhibited. |
| Fig. 4g | 41A01>TNT, tsh-Gal80 | Courtship-promoting pCd1 neurons were constitutively silenced in *Dmel* brain. |
| Fig. 4g | 41A01> Kir, tub-Gal80$^{TS}$ | pCd1 neurons were conditionally silenced in adult *Dmel* flies. |
| Fig. 4h | 41A01>GFP | pCd1 neurons were labeled with GFP in the *Dmel* brain. |
| Fig. 4i | TH>P2X$_2$; 41A01>GCaMP6s | Genetic line where dopamine neurons were conditionally activated through exogenous ATP application, and neural calcium activity was simultaneously recorded from pCd1 neurons. |
| Fig. 4j | 41A01>RetroTango | Genetic line to label neurons pre-synaptic to pCd1 neurons. |
| Fig. 4k | 41A01>Dop1R2$^{KO}$ | The dopamine receptor Dop1R2 was selectively removed from pCd1 neurons using CRISPR. |
| Fig. 4k | 57C10>Dop1R2$^{KO}$ | The dopamine receptor Dop1R2 was removed from all neurons of *Dmel* central nervous system using CRISPR. |
| ED Fig. 4c | nsyb>NanRNAi | Mechanosensory ion channel Nanchung was knocked down in all neurons of *Dmel* central nervous system. |
| ED Fig. 4f | ppk23>hid | CHC sensing ppk23 neurons were ablated in *Dmel* males. |
| ED Fig. 4f | ppk23>Kir, tub-Gal80$^{TS}$ | ppk23 neurons were conditionally silenced in adult *Dmel* flies. |
| ED Fig. 5g | Oe$^{-}$ | Genetic line where oenocytes, the pheromone-producing cells, were ablated, rendering the flies pheromone-deficient. |
| ED Fig. 7a | DAT>TNT, tsh-Gal80 | Activity of all dopaminergic neurons in the *Dmel* brain was constitutively silenced. |
| ED Fig. 7a | DAT>Kir, tub-Gal80$^{TS}$ | All dopamine neurons were conditionally silenced in the adult *Dmel* flies. |

| | | |
|---|---|---|
| ED Fig. 7h | TH-F2>TNT, tsh-Gal80; TH-D4>TNT, tsh-Gal80 | Activities of dopamine neuron subsets (TH-F2, TH-D4) were silenced. |
| ED Fig. 7j | 58E02>TNT, tsh-Gal80; 92G05>TNT, tsh-Gal80 | Neural activities of the dopamine cluster PAM and PPM3 were silenced. |
| ED Fig. 8b | Crz>NaChBac | Corazonin neurons were hyperactivated in *Dmel* males. |
| ED Fig. 9a | 69F08>CaLexA | Calcium-dependent reporter was expressed to mark the activation of sleep-regulating ellipsoid body (EB) neurons. |
| ED Fig. 9b | 23E10>CaLexA | Calcium-dependent reporter was expressed to mark the activation of sleep-promoting dorsal fan-shaped body (dFB) neurons. |
| ED Fig. 9f | PPM2sp>NaChBac | PPM2 dopamine neurons were constitutively hyperactivated in *Dmel* males. |
| ED Fig. 9f | PPM2sp>TrpA1 | PPM2 neurons were conditionally hyperactivated in adult *Dmel* males. |
| ED Fig. 9g | 23E10>NaChBac | Sleep-promoting dFB neurons were hyperactivated in *Dmel* males. |

**Supplementary Table 4:** Behavioral features associated with HMM-inferred states.

| | Interaction | Ambulation | Micro movements | Feeding | Grooming | Rest |
|---|---|---|---|---|---|---|
| **State 1** | Moderate | Moderate | Low | Moderate | Moderate | High |
| **State 2** | High | Low | High | High | Low | Moderate |
| **State 3** | Moderate | High | Moderate | Low | High | Low |
