## supplementary table 1 for "Mating imperatives drive plasticity of the daily temporal niche in fruit flies"

| *Dmel* **wild strains** | **Collected from** | **Approx. Coordinate** |
|---|---|---|
| *D. melanogaster* Acc1 | Accra, Ghana | 5.3°N/0°E |
| *D. melanogaster* Oku | Oku, Cameroon | 6.2°N/10.5°E |
| *D. melanogaster* Tan | Machame, Tanzania | 3.11°S/37.14°E |
| *D. melanogaster* Zam | Zambia | 15.5°S/28.3°E |
| *D. melanogaster* Zim S-29 | Zimbabwe | 18.08°S/28.19°E |
| *D. melanogaster* Mal 63 | Malawi | N/A |
| *D. melanogaster* Hub11 | Bayern, Germany | 49.8°N/9.9°E |
| *D. melanogaster* Kop3 | Copenhagen, Denmark | 56.2°N/12.6°E |
| *D. melanogaster* Lei38 | Leicestershire, England | 52.6°N/1.1°W |
| *D. melanogaster* Vil | Viltain, Saclay, France | 48.76°N/2.15°E |
| *D. melanogaster* Ban | Bangalore, India | 12.80°N/77.58°E |
| *D. melanogaster* Bre | Bretagne, France | 48.06°N/3.08°W |
